# Spatial Transcriptomics of Intraductal Papillary Mucinous Neoplasms of The Pancreas Identifies NKX6-2 as a Driver of Gastric Differentiation and Indolent Biological Potential

**DOI:** 10.1101/2022.10.19.512773

**Authors:** Marta Sans, Yuki Makino, Jimin Min, Kimal I. Rajapakshe, Michele Yip-Schneider, C Max Schmidt, Mark W. Hurd, Jared K. Burks, Javier A. Gomez, Fredrik I. Thege, Johannes Fahrmann, Robert A. Wolff, Michael P. Kim, Paola A. Guerrero, Anirban Maitra

## Abstract

Intraductal Papillary Mucinous Neoplasms (IPMNs) of the pancreas are *bona fide* precursor lesions of pancreatic ductal adenocarcinoma (PDAC). The most common subtype of IPMNs harbor a gastric foveolar-type epithelium, and these low-grade mucinous neoplasms are harbingers of IPMNs with high-grade dysplasia and cancer. The molecular underpinning of gastric differentiation in IPMNs is unknown, although identifying drivers of this indolent phenotype might enable opportunities for intercepting progression to high-grade IPMN and cancer. We conducted spatial transcriptomics on a cohort of IPMNs, followed by orthogonal and cross species validation studies, which established the transcription factor NKX6-2 as a key determinant of gastric cell identity in low-grade IPMNs. Loss of *NKX6-2* expression is a consistent feature of IPMN progression, while re-expression of *NKX6-2* in murine IPMN lines recapitulates the aforementioned gastric transcriptional program and glandular morphology. Our study identifies NKX6-2 as a previously unknown transcription factor driving indolent gastric differentiation in IPMN pathogenesis.

**Significance:** Improved understanding of the molecular features driving IPMN development and differentiation is critical to prevent cancer progression and enhance risk stratification. Our study employed spatial profiling technologies to characterize the epithelium and microenvironment of IPMN, which revealed a previously unknown link between *NKX6-2* expression and gastric differentiation in IPMNs, the latter associated with an indolent biological potential.

## Introduction

Pancreatic Ductal Adenocarcinoma (PDAC) is projected to become the second leading cause of cancer death by the year 2040 (1). Significantly higher survival rates are observed in patients diagnosed with localized (42%) compared to regional or distant disease (14% and 3%, respectively), emphasizing the need for early detection and interception strategies to improve patient outcomes (2). Intraductal Papillary Mucinous Neoplasms (IPMNs) are the most common category of mucin-producing pancreatic cysts and can arise from either the main pancreatic duct, or from one or more of the branch ducts. IPMNs are *bona fide* precursor lesions of PDAC, with ∼5-10% of non-invasive cystic lesions progressing to invasive adenocarcinoma (3,4). Based on the degree of atypia, the lining epithelium of IPMNs is classified into either low-grade (LG) or high-grade (HG) dysplasia, with the latter harboring a greater propensity for progression to invasive neoplasia (5). In addition, based on morphological features and parallels to the epithelial cells from other gastrointestinal organs, the lining epithelium of IPMNs has also been categorized into gastric, intestinal, or pancreato-biliary subtypes (6). The gastric subtype is by far the most common subtype of IPMNs, mostly arising within the branch ducts, and is typically associated with LG dysplasia. In contrast, IPMNs of the intestinal and pancreato-biliary subtypes can arise in either the main or branch ducts, and usually correspond to HG IPMNs. The phylogeny of intestinal and pancreato-biliary subtypes continues to be debated, with some studies speculating that gastric IPMNs represent a common precursor to both categories of HG lesions (7,8), and others postulating that gastric and pancreato-biliary IPMNs share a developmental trajectory that is distinct from the intestinal subtype (9,10). Irrespective, the deregulated transcriptional programs that characterize the most common category of IPMNs, the gastric subtype, remain mostly an enigma. Identifying drivers of gastric differentiation within the IPMN epithelium would not only facilitate insights into the pathogenesis of these early cystic lesions but would also provide the basis for cataloging alterations that accompany further progression to HG IPMNs, and eventually cancer.

Newly emergent spatial technologies for transcriptome and protein profiling offer the opportunity to evaluate the levels of gene transcripts and proteins directly from tissue samples while preserving their localization and distribution (11,12). In this study, we utilized the Visium spatial transcriptomics (ST) platform to profile snap-frozen archival samples from a panel of patient IPMNs with varying grades of dysplasia and epithelial morphology. With this approach, we identified distinct epithelial patterns of gene expression that were associated with IPMN grade and subtype. Using single-cell deconvolution algorithms, the cellular composition of the corresponding IPMN-associated microenvironment was also characterized. Notably, we identified that elevated levels of transcripts encoding the transcription factor *NKX6-2* (NK6 homeobox 2) were localized to the epithelium of LG IPMNs and were associated with a gene signature recapitulating the gastric pit and isthmus epithelium, which we have termed gastric isthmus-pit (GIP) signature. We validated the GIP signature associated with elevated *NKX6-2* expression using multiple orthogonal approaches, including RNA sequencing performed on archival microdissected IPMN samples and cross-species spatial transcriptomics performed in a genetically engineered model of IPMNs (13,14). Moreover, using a quantitative high-plex platform for protein profiling on archival material (Lunaphore COMET assay), we confirmed that elevated nuclear NKX6-2 was a feature of the gastric subtype of IPMNs, wherein this transcription was co-localized with markers of gastric foveolar mucin (TFF1 and MUC5AC), and with nuclear GATA6, a prototypal marker of so-called “classical” differentiation within pancreatic neoplasms (15). The GIP signature was also recapitulated with overexpression of NKX6-2 in cells derived from a genetically engineered model of IPMNs. Remarkably, orthotopic implantation of *NKX6-2* overexpressing murine IPMN-derived cancer cells resulted in lesions with substantial glandular differentiation recapitulating IPMN-like lesions, compared to the predominantly undifferentiated parental cells. Cumulatively, our data support a hitherto unknown role for NKX6-2 in driving a gastric-type transcriptional profile in the most commonly observed subtype of IPMNs, and a master regulator of the glandular differentiation that characterizes these lesions.

## Results

### Spatially resolved cell-type specific expression profiles of IPMN samples

The archival snap-frozen IPMN cohort analyzed in our series was comprised of seven LG IPMNs, three HG IPMNs, and three IPMN-associated cancers (**Table S1**). Of note, all LG cases in the dataset were categorized as gastric subtype, while HG cases included pancreato-biliary and intestinal subtypes of IPMN. ST analyses on the thirteen specimens yielded spatially resolved expression profiles for up to 8,000 genes per each of the 12,685 spots covering the tissue samples (**Fig. S1**). Utilizing the scMC (16) processing and clustering pipeline, spots were grouped into 14 clusters according to their gene expression profiles (**Fig. 1A, Table S2**). The majority of clusters included spots from a multitude of samples, while others were predominantly contributed by one or two samples (**Fig. S2**). Visualization of the spatial localization of the spots for each of the clusters showed correlations with histologic al features. Cell-type proportions for each of the spots were predicted using the robust cell type deconvolution (RCTD) algorithm (17), with the overall highest proportions per spot attributed to epithelial cells and fibroblasts. Not unexpectedly, spots enriched in epithelial cells corresponded to IPMN epithelium while spots containing fibroblasts were in areas of high stromal density (**Table S3**). On the other hand, most immune cell types garnered lower percentage scores per spot, which was expected due to the overall sparsity of such cell types within the IPMN microenvironment. As shown in **Fig. 1B**, trends in cell composition were observed for each of the clusters, which also aligned with the histological assignments. For example, clusters 7 and 11 included the highest average epithelial cell percentages, expressing elevated levels of *EPCAM* transcripts, and were found to be localized to the epithelial cells in the IPMN and associated cancer samples (**Fig. 1C**). High proportion of fibroblasts were found in clusters 3, 9, 13 and 14, which corresponded to stromal tissue areas with high expression of collagen genes (*COL1A1, COL3A1, COL1A2*). Spots located in tissue areas containing lymphocyte aggregates in cluster 2 were comprised of the highest per spot percentages in naïve lymphocyte cells and expressed elevated levels of T and B cell markers, such as *CR2* and *CD52* (**Fig. 1D**). High percentage of plasma cells (up to 65%) were found in clusters 4 and 6, which were derived from the stroma adjacent to IPMN epithelium and showed elevated expression of various immunoglobulin transcripts (i.e., *IGHA1, IGLC2*) (**Fig. 1E**). Overall, unsupervised characterization of the transcriptional profiles obtained from ST analyses from whole tissue samples, together with deconvolution approaches, provided a correlation between gene expression and tissue histology.

**Figure 1.**
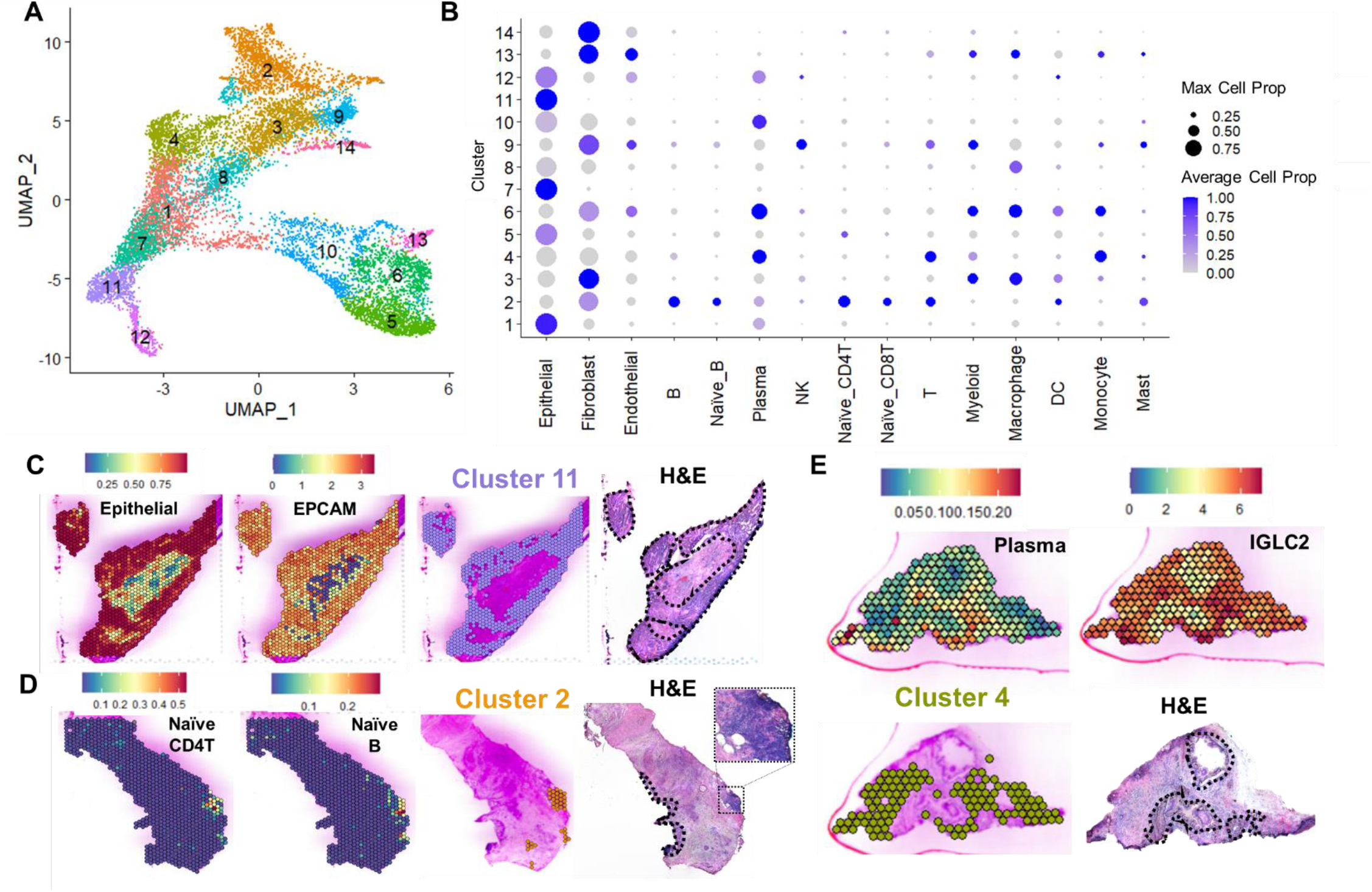
Spatially resolved transcriptomic analyses of IPMN tissue samples. A) Uniform Manifold Approximation and Projection (UMAP) showing clustering of spots based on gene expression profiles for all 13 samples. B) Dot plot including cell type scores per cluster obtained with the robust cell type decomposition (RCTD) algorithm. Color represents the normalized average cell proportion and dot size represents the maximum cell proportion per cluster. Representative spatial maps showing correlation between cell composition and localization provided by RCTD, spatial gene expression, cluster assignments, and histology. C) Mapping of epithelial cells, EPCAM expression, and localization of spots from cluster 11 for a HG IPMN sample. D) Localization and proportion of naïve CD4T and naïve B cells, and visualization of cluster 2 spots from a HG IPMN sample with co-occurring cancer. A tissue area corresponding to a lymphocyte aggregate is outlined and a zoomed-in image is provided. (E) Plasma cells, IGLC2 expression, and cluster 4 shown for a LG IPMN sample. IPMN areas are outlined in black in the optical images from H&E stained tissues.

### Region-specific differential expression of transcripts and cell types associated with IPMN progression

Clinically, HG IPMNs are considered as high-risk lesions that mandate surgical resection, with or without an associated invasive cancer. Therefore, for purposes of further analyses, the three HG IPMNs and three IPMN-associated cancers in our ST cohort were combined into one “high-risk” (HR) category for comparison *versus* the LG IPMNs. To evaluate region-specific changes in gene expression and cell composition between the LG IPMNs and HR IPMNs, we selected spots overlapping with the neoplastic epithelium (“epilesional”, n=755); the immediately adjacent microenvironment, corresponding to two layers of spots (∼200 µm) surrounding the lining epithelium (“juxtalesional”, n=1142); and an additional two layers of spots located further distal to the juxtalesional region (“perilesional”, n=1030) (**Fig. 2A**). Comparisons between the transcriptomic profiles of LG IPMN *versus* HR IPMNs resulted in the identification of 946, 705, 272 transcripts that were significantly differentially expressed in the epilesional, juxtalesional, and perilesional areas, respectively, between the two categories of IPMNs (**Table S4-S6, Fig. 2B-D**). Differential expression analysis was also conducted between the regional compartments from each of the three tissue types (LG, HG, and PDAC) (**Table S7**). Interestingly, expression of genes associated with gastric mucus-secreting cells, such as *MUC6, NKX6-2, PGC, PSCA*, and *TFF2* were found to be elevated in the LG IPMN epithelium when compared to HR IPMN. Similarly, higher expression levels of *IGLC2* and *IGKC*, markers of activated B cells and plasma cells, were detected in the LG IPMN epilesional areas, suggesting infiltration of plasma cells in these lesions. On the other hand, transcripts previously described as upregulated in “conventional” PDAC, such as *CEACAM6* and *CLDN4* were found to be expressed at higher levels in HR IPMN samples when compared to LG IPMN (18,19). Transcripts with higher expression restricted to HG IPMN compared to either LG IPMN or PDAC epilesional areas were associated with intestinal differentiation, such as *CDX1, CDX2, MUC2*, and *CDH17*, and were largely driven by one HG intestinal type IPMN in our dataset (**Fig. S3A-B**). Not unexpectedly, these features were also observed in an invasive “colloid” carcinoma, which arose in the backdrop of an intestinal IPMN. Higher expression of interferon stimulated genes, such as *BST2, ISG15*, and *LY6E*, and of *KLF7*, which has been implicated in promoting cancer growth by up-regulating interferon induced genes (20) were observed at higher levels in PDAC epilesional spots compared to both LG and HG IPMN (**Fig. S3C**)

**Figure 2.**
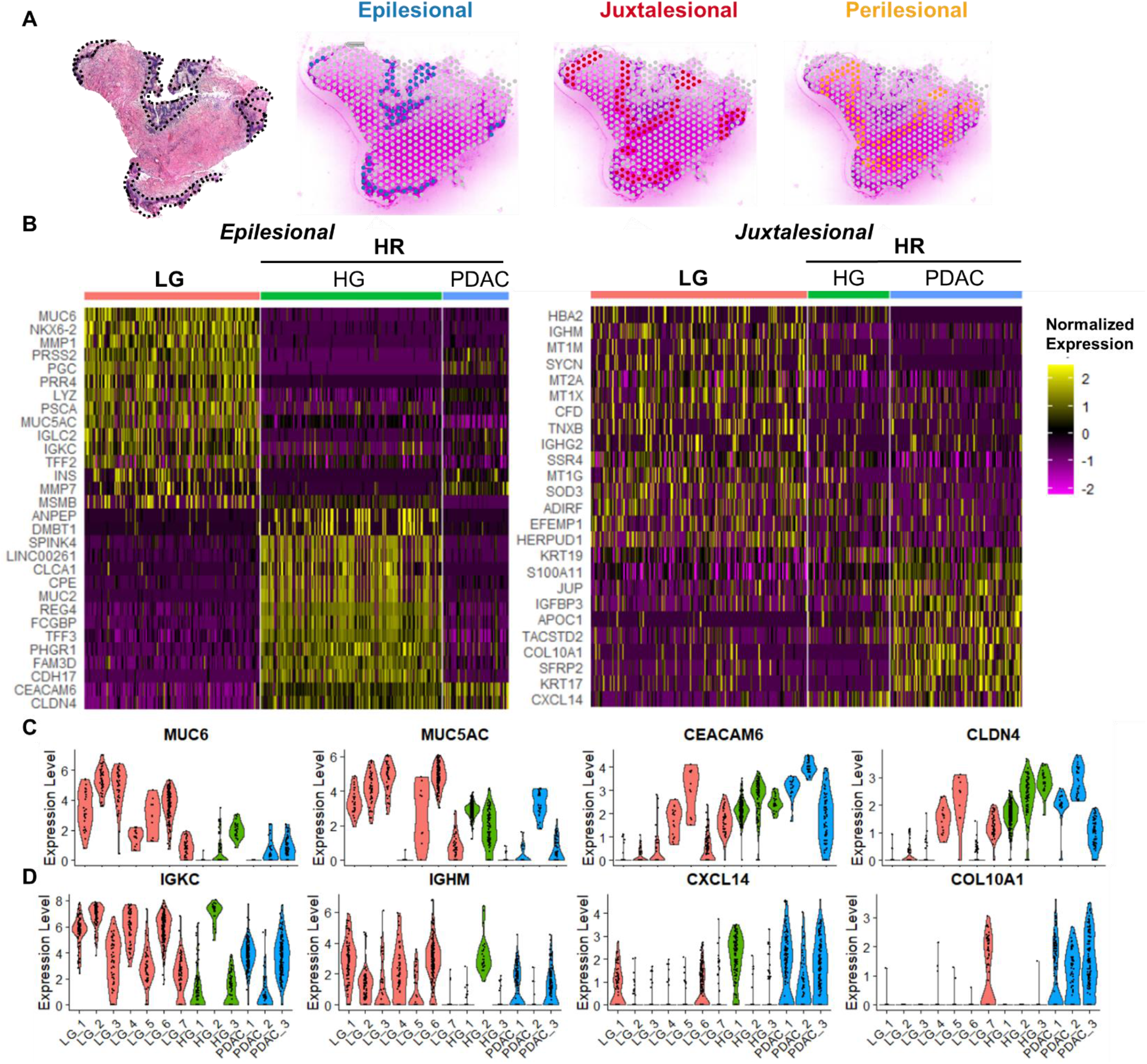
Histologically directed spot selection reveals area-specific gene expression changes associated with IPMN grade and PDAC. A) Selection of epilesional, juxtalesional, and perilesional areas shown in a LG IPMN sample. B) Heatmaps plotting expression of genes differentially expressed between LG and HR (high-risk) in epilesional (left) and juxtalesional (right) spots. Transcripts exclusively showing differential expression in the jutxalesional and not the epilesional compartments were included in the juxtalesional heatmap. Violin plot showing gene expression levels for selected features for spots in C) epilesional and D) juxtalesional areas.

In the juxtalesional region, higher levels of various immunoglobulin transcripts, including those also observed in the epilesional areas (*IGLC2, IGKC*) and others unique to this compartment (*IGHM, IGHG2*) were observed in LG IPMN samples compared to HR IPMNs, reiterating the role of an active humoral immune response in early *versus* late-stage cystic lesions. Transcripts corresponding to cancer associated fibroblasts, like *COL10A1*, were found to be significantly higher expressed in the juxtalesional regions surrounding HR IPMNs, and in particular, PDAC samples, compared to those surrounding LG IPMN. A lower number of differentially expressed genes were present when comparing the perilesional areas, suggesting that changes in the microenvironment between LG and HR IPMN are more prominent in the areas closest to the lesions.

Utilizing the cell-type proportions assigned by RCTD, we then evaluated the cell composition in the three selected compartments in our IPMN cohort (**Fig. 3A**). Trends in cell proportions were observed when comparing the three regions between LG IPMN *versus* HR IPMN, even though large variance was observed between spots and patients, suggesting heterogeneous spatial distribution of cell types (**Table S8, Fig. S4A**). Of note, significantly higher proportions of plasma and mast cells were assigned in the jutxalesional areas of LG IPMN compared to HR IPMNs (**Fig. 3B**). By RCTD, juxtalesional spots were predicted to contain an average of 10% plasma cells in LG IPMN cells, while an average of 5% was observed in HR IPMN samples, with a single case (HG_2) contributing to most of the plasma cell counts (**Fig. 3C**). Spots with high levels of plasma cells were also characterized by expression of immunoglobulin genes, such as *IGKC* (**Fig. S4B**). Mast cells were predicted to comprise as much as 10% of the cells in the juxtalesional spots, suggesting a significant infiltration of such cells in the IPMN microenvironment (**Fig. 3B, D**). Spots with mast cell contributions showed high levels of *TPSAB1*, a transcript encoding for mast-cell specific tryptase (**Fig. S4C**).

**Figure 3.**
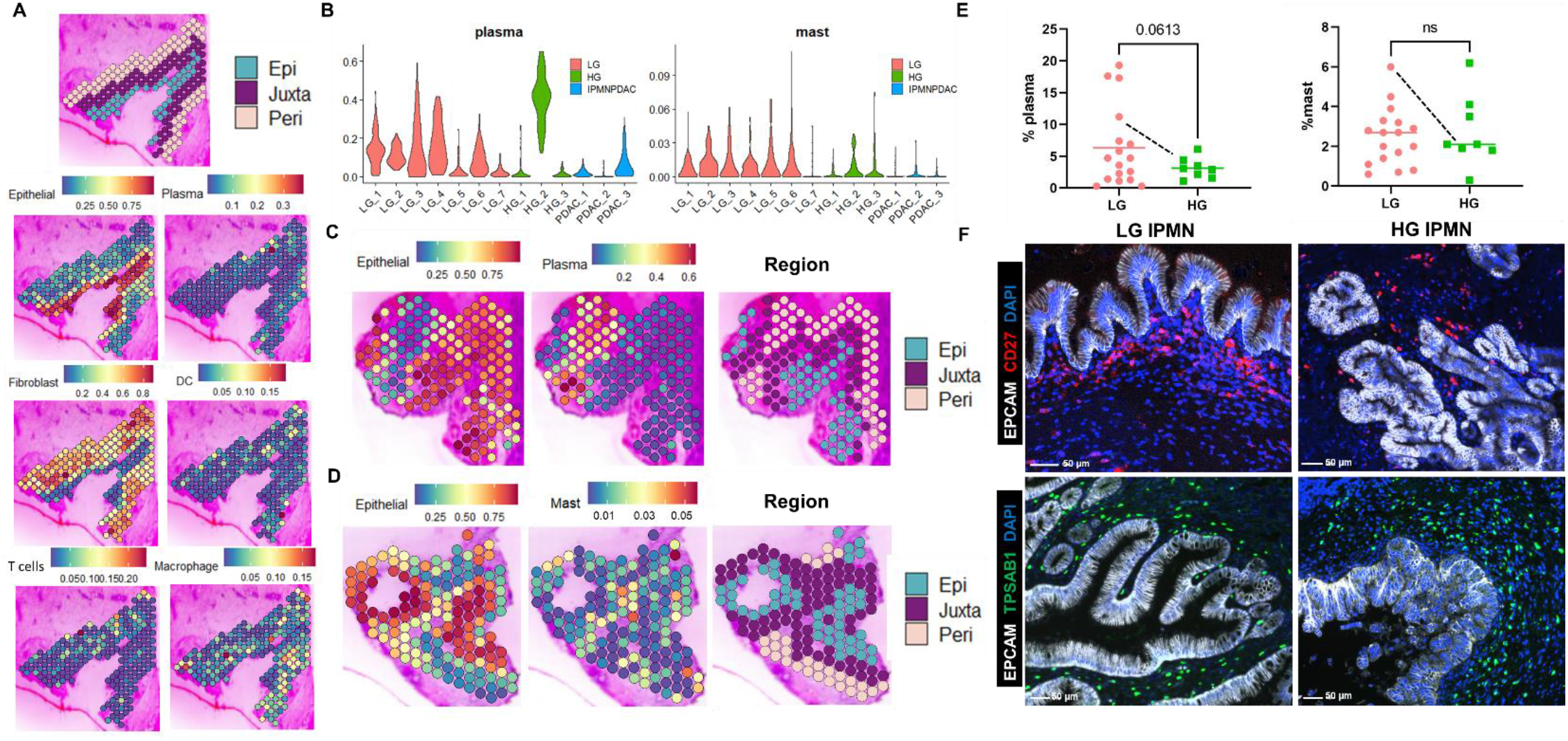
Spatial organization of the tumor microenvironment in IPMN samples. A) Representative spatial maps showing proportion of cell types in the ST spots as predicted by RCTD. The image at the top shows the spots colored according to their region. B) Violin plot showing per spot proportions of plasma and mast cells grouped by sample code and colored by type. Representative spatial maps highlighting the proportion and localization of epithelial, plasma (C), and mast (D) cells in the IPMN samples. On the right side, spots are colored according to their corresponding region (epilesional, jutxalesional, and perilesional). E) Percentage of cells staining positive for CD27 (left) and TPSAB1(right) in the adjacent microenvironment to the IPMN lesions in the samples analyzed by the COMET imaging system. A dash line connects the LG and HG IPMN lesions that correspond to the same patient and tissue section. F) COMET images showing co-staining of CD27 (top) and TPSAB1 (bottom) with EPCAM and DAPI in LG and HG IPMN samples.

To further evaluate the presence of plasma and mast cells in the IPMN microenvironment, we quantified the expression of CD27 (plasma cells) and TPSAB1 (mast cells) in the IPMN-adjacent microenvironment by cyclic IF (COMET) in an independent cohort of archival formal-fixed paraffin embedded (FFPE) IPMNs (**Fig. 3E-F, Table S9**). The COMET analysis confirmed that a higher percentage of plasma cells was observed in LG *versus* HG IPMN, which albeit not reaching statistical significance, demonstrated a similar trend to that observed on the ST data. In contrast, mast cell percentages in the adjacent microenvironment from both LG and HG IPMNs were comparable (**Table S10**). Nonetheless, when comparing the adjacent microenvironment between synchronous LG and HG IPMN areas in the same tissue section, a higher percentage of both plasma and mast cells was observed surrounding the LG (11.2% and 4.4%, respectively) compared to the HG lesions (6.0% and 1.1%, respectively) (**Fig. 3E, Fig. S5A**). Utilizing the additional markers in the COMET panel, other immune cell phenotypes, including macrophages, T cells, and B cells were quantified in the adjacent microenvironment. Although we observed trends between LG and HG IPMN samples with similar directionality as ST analysis, none of these comparisons were found to be statistically significant (**Table S10, Fig. S5B**). Thus, given the broad heterogeneity in cell type composition of the juxtalesional microenvironment observed in the IPMN lesions, which has been previously described by others (21), we decided to focus on further dissecting the epilesional compartment of LG and HG IPMN lesions to identify potential markers of grade and subtype.

### Elevated epithelial NKX6-2 expression is associated with gastric differentiation in IPMNs

While the gastric subtype is the most commonly observed variant of IPMNs in the pancreas, the putative transcriptional drivers underlying this pattern of differentiation are not known. In our ST dataset, all LG IPMNs were uniformly comprised of the gastric subtype, while HG IPMNs were either intestinal or pancreato-biliary, and thus the terms “LG” and “gastric” are used interchangeably. As previously noted, expression of genes associated with gastric mucus-secreting cells were found to be elevated in the epilesional regions of LG IPMN *versus* HR IPMNs (**Fig. 2B-C**). Of these, *MUC6* was the highest differentially expressed transcript, which was not surprising given the encoded apomucin is a prototype gastric mucin. The second highest differentially expressed gene in the epilesional regions of LG IPMNs versus HR IPMNs was *NKX6-2* (**Fig. 2B, Fig. 4A-B**), which encodes for a homeobox domain containing transcription factor. *NKX6-2* is expressed in pancreatic endocrine cells, but not in normal ductal cells, and is known to promote beta-cell differentiation (22,23). *NKX6-2* is also highly expressed in human stomach, particularly in isthmus and pit cells (24,25). Moreover, a homolog to *NKX6-2, Nkx2-1*, has been implicated in regulating gastric differentiation in the lung (26). To validate NKX6-2 expression in IPMN, we performed quantitative analysis of nuclear NKX6-2 expression using the COMET platform on an independent cohort of archival IPMN samples, which also demonstrated a significantly higher percentage of nuclei staining positive for NKX6-2 in LG (all gastric subtype) compared to HG IPMN samples, which included both intestinal and pancreato-biliary subtypes (**Fig. 4C-D, Table S9, Table S11**). Notably, in a case with synchronous LG and HG IPMN foci in the same section, a higher proportion of NKX6-2-expressing nuclei were observed in the LG nuclei (25%) *versus* HG IPMN (1%) (**Fig. 4E**). Agreements between *NKX6-2* levels determined by ST and the percent of nuclei positive for NKX6-2 in the COMET images was also observed for the single patient with tissue analyzed by both methods (**Fig. S6**). Despite the overall significantly higher levels of expression of this transcription factor in LG IPMNs compared to HG IPMNs, we nonetheless observed inter- and intra-patient heterogeneity in expression of both *NKX6-2* transcripts and protein within the LG IPMN epithelium. In instances of reduced expression of nuclear NKX6-2 in LG IPMN, this pattern was usually localized to transitory states from gastric to pancreato-biliary IPMN, underscoring that loss of this transcription factor might be an accompaniment of progression (**Fig. S6**).

**Figure 4.**
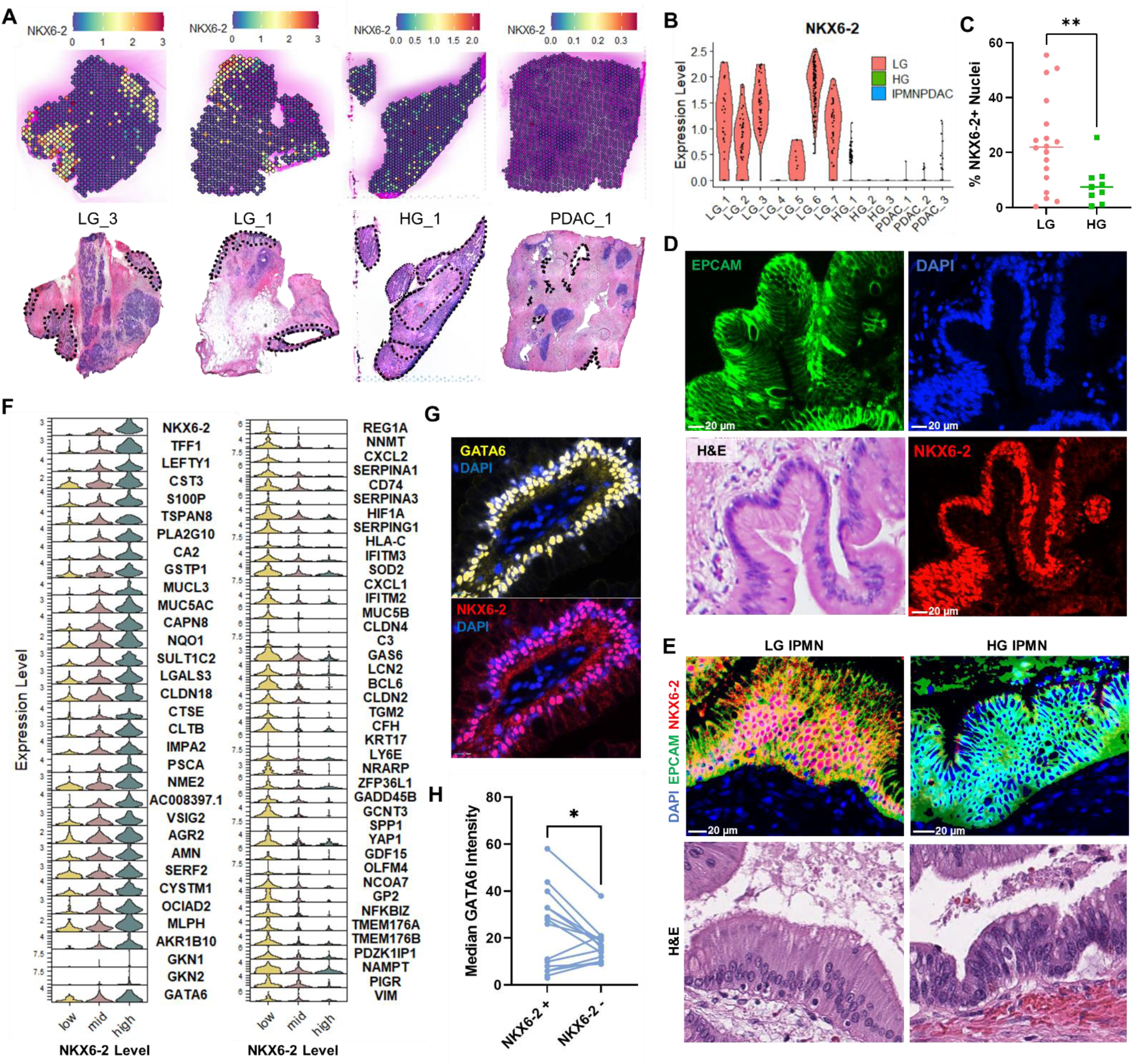
*NKX6-2* expression is elevated in LG IPMN and associated with a distinct transcriptional profile. A) Spatial maps of *NKX6-2* expression and localization from ST analyses of four representative samples. Optical images for the corresponding H&E images are provided below, with the neoplastic epithelium outlined. B) Violin plot showing *NKX6-2* expression in the IPMN epilesional spots grouped by sample. C) Percentage of NKX6-2 positive nuclei in IPMN epithelium determined by COMET cyclic IF staining (p = 0.0039). D) Localization of NKX6-2 in the nuclei of LG IPMN epithelium showed by co-staining with EPCAM, DAPI, and NKX6-2. An H&E image of the area stained is provided in the bottom left. E) NKX6-2 staining of LG and HG IPMN lesions from the same tissue section showing higher NKX6-2 levels in the LG IPMN nuclei. H&E images of the areas stained are provided below. F) Representative gene transcripts observed at significantly higher levels in spots of high *NKX6-2* (left panel) or low *NKX6-2* levels (right panel) in the ST dataset from LG IPMN. G) Co-staining of GATA6 and NKX6-2 showing co-expression in the LG IPMN nuclei. H) Median GATA6 intensity obtained from the COMET staining in NKX6-2+ versus NKX6-2- nuclei (p = 0.03).

To interrogate molecular changes associated with *NKX6-2* expression, epilesional spots in the ST dataset corresponding to the LG IPMN epithelium were separated based on *NKX6-2* levels into “low”, “mid”, and “high”, and differential expression analyses were performed (**Fig. S7A-B, Table S12**). Importantly, the three categories included spots from almost all samples, allowing us to perform both inter and intra-patient comparisons. Higher levels of transcripts comprising a gastric isthmus-pit (GIP) signature were detected in epilesional spots of high and mid versus low NKX6-2 spots, including *TFF1, LEFTY1, CST3, S100P, TSPAN8, PLA2G10, MUC5AC*, and *GKN2*, among others (**Fig. 4F, S7C**). On the other hand, higher expression of *CXCL2, CD74, CLDN4, LY6E, and SPP1*, many of which have been associated with PDAC progression and neoplastic progression of IPMN (18,27-29), was observed in epilesional spots with minimal *NKX6-2* expression (**Fig. 4F, S7D)**. Expression of transcripts associated with the classical subtype of PDAC, such as *GATA6*, was also higher in epilesional spots with high *NKX6-2* expression (15,30). Moreover, GATA6 and NKX6-2 protein co-localization was observed in the nuclei of LG IPMN using the COMET platform, with overall significantly higher median intensity of GATA6 observed in NKX6-2 positive versus negative nuclei (**Fig. 4G-H, Table S13**). On the contrary, LG epilesional spots with low *NKX6-2* expression had lower levels of classical-like genes, and a percentage of those (∼25%) expressed prototypal markers of the basal subtype, including *KRT17 (31)* (**Fig. S7E**). Higher levels of *KRT17* have also been observed in pancreato-biliary, compared to both intestinal and gastric IPMNs (10). Of note, acquisition of intestinal IPMN markers, *CDX2* or *MUC2*, was not observed in the LG gastric IPMN lesions upon *NKX6-2* loss, thus not supporting a transition from gastric to intestinal subtype in these cases. Additionally, gene set enrichment analysis (GSEA) revealed increased expression of genes associated with epithelial-to-mesenchymal transition (EMT) in areas with loss of NKX6-2 (**Fig. S7F-G**).

Additional GSEA analyses using cell type signatures showed enrichment of gene sets obtained from gastric pit and isthmus cells from previously published single cell RNA sequencing datasets of human normal stomach (**Fig. 5A, Table S14**) in epilesional spots expressing *NKX6-2* (high and mid) compared to spots with low expression in the LG IPMN ST dataset (25). These results further corroborated the association of *NKX6-2* expression with the GIP signature. Gene sets corresponding to gastric REG3A positive cells, fetal pancreas acinar cells, and pancreas ductal cells were enriched in the transcriptional profiles obtained from spots with minimal NKX6-2 expression (**Fig. S8A**). To further evaluate tissue level expression of NKX6-2 together with markers of the GIP signature, we interrogated TFF1 and MUC5AC co-expression with NKX6-2 using the COMET cyclic IF panel, and observed localization of both markers in the cytoplasm of LG IPMN epithelium, preferentially in areas with NKX6-2 positively stained nuclei (**Fig. 5B**). Quantification of per sample protein expression in the IPMN epithelium showed moderately strong positive correlations between percentage of positive NKX6-2 nuclei with MUC5AC (cor = 0.54, p-val = 0.02) and TFF1 mean intensities (cor = 0.55, p-val = 0.02) (**Table S13, Fig. S8B**).

**Figure 5.**
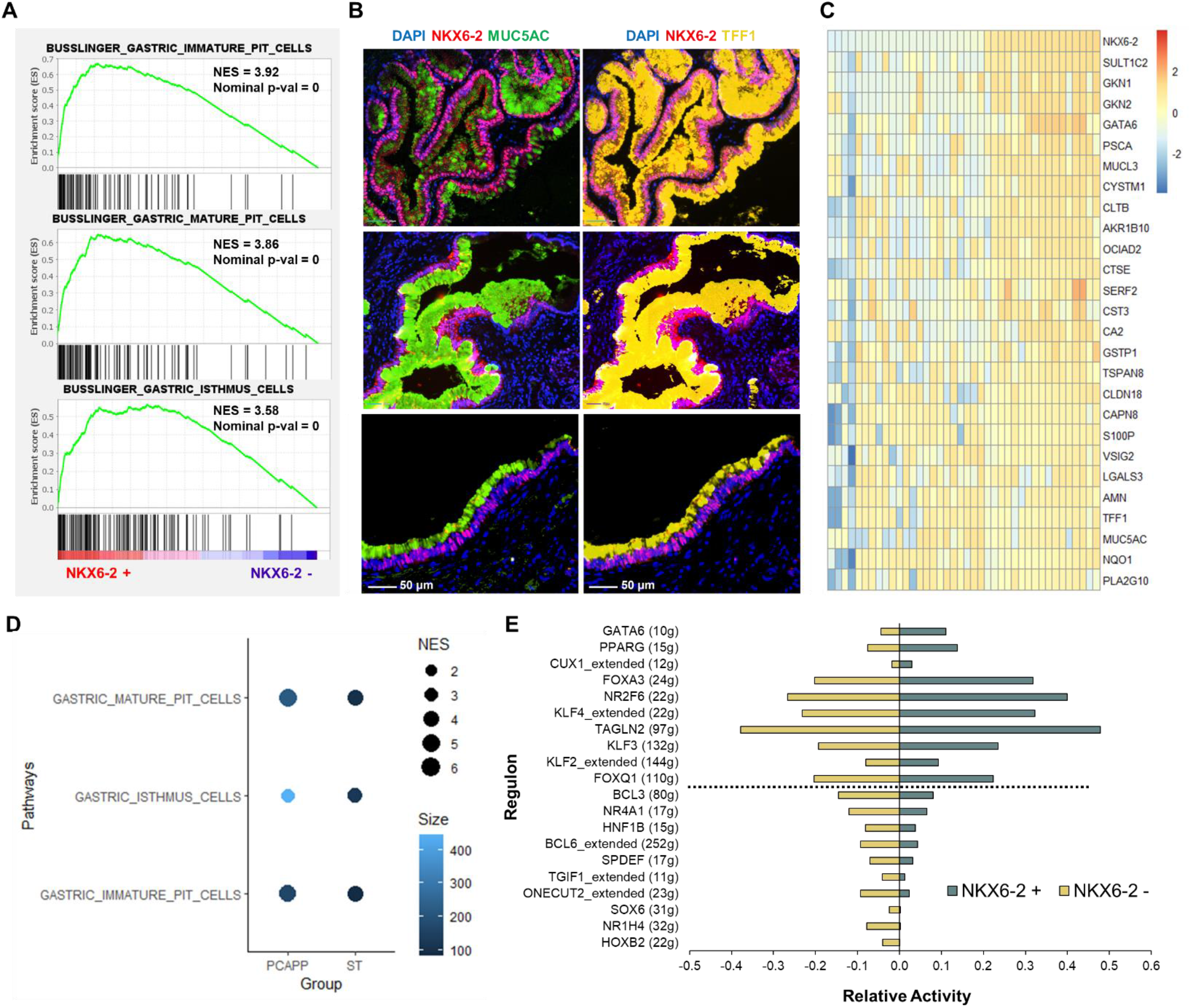
Enrichment of the GIP signature in *NKX6-2* expressing IPMN. A) Enrichment of gastric pit and isthmus gene sets in the *NKX6-2* associated signature obtained from the ST LG IPMN dataset. B) Representative COMET images showing co-staining of NKX6-2, MUC5AC, and TFF1 in three different LG IPMN samples. C) Heatmap showing expression of *NKX6-2* and the GIP signature in 23 archival IPMN tissue areas from the NCI Cancer Moonshot PCAPP dataset (14). Samples are ranked by *NKX6-2* expression from right to left. Heatmap values represent normalized RNA-seq expression values. D) Dot plot showing normalized enrichment scores and size for gastric pit and isthmus gene sets in both the PCAPP and ST dataset by GSEA. E) Regulon activities scored for *NKX6-2*+ (green) and *NKX6-2*- (light yellow) spots based on SCENIC analyses.

To validate whether *NKX6-2* expression in IPMNs correlates with a gastric transcriptional program, we analyzed an independent dataset of RNA sequencing obtained following laser-capture microdissection (LCM) of 23 archival IPMNs, conducted as part of the NCI Cancer Moonshot PreCancer Atlas Pilot Project (PCAPP) (14). Remarkably, majority of the transcripts that comprise the aforementioned GIP signature (including *TFF1, GKN1, GKN2*, and *MUC5AC*, amongst several others), as well as the prototypal classical-like transcript *GATA6* were also positively correlated with *NKX6-2* expression in the PCAPP dataset (**Fig. 5C, Fig. S8C**). GSEA also revealed enrichment of the gastric pit and isthmus gene sets with *NKX6-2* expression in the PCAPP cohort similar to what was observed in the ST dataset (**Fig. 5D**). Thus, in two independent datasets of archival IPMNs profiled by disparate approaches (ST and RNA-sequencing), elevated *NKX6-2* was consistently associated with the GIP signature.

### Status of NKX6-2 expression in IPMNs correlates with alterations in regulatory networks

We applied SCENIC (32,33) to evaluate regulatory networks that might be associated with *NKX6-2* expression utilizing the ST dataset. Using this approach, 6,266 genes were identified as targets of the *NKX6-2* transcription factor. The targets identified included 119 out of the 134 genes selected as significantly higher expressed in *NKX6-2* positive *versus NKX6-2* negative epilesional spots in LG IPMN from the ST analyses, with markers of the GIP signature, including *TFF1, CAPN8, CA2, SULT1C2, S100P*, and *PLA2G10*, selected among the targets receiving the highest weights (**Table S15**). We then applied *RcisTarget* to identify “regulons”, defined as gene modules that are co-regulated by a main transcription factor (**Table S16**). We identified the GATA6 regulon presenting the highest positive fold change in activity between *NKX6-2* positive versus *NKX6-2* negative spots, reiterating the link between NKX6-2 and classical differentiation. *FOXA3* was among the other regulons with the highest score positively correlated with NKX6-2 expression (**Fig. 5E**). Higher levels of the *FOXA3* transcriptional network have been reported in the pancreatic progenitor subtype of PDAC, which expresses a foregut-related differentiation program, compared to the more mesenchymal/squamous or basal subtypes (34). Interestingly, Foxa1 and Foxa2, homologs to Foxa3, have been found to physically interact with lineage specific homeobox transcription factors, such as Nkx2-1 in the pulmonary epithelium, to drive a gastric differentiation program in lung carcinomas (26,35). *KLF*-related regulons, such as *KLF4, KLF3*, and *KLF2*, were also observed at higher activities in *NKX6-2* expressing spots. Gastric epithelial proliferation and differentiation, including that of pit cells, is largely controlled by *KLF4*, which provides essential functions to regulate normal gastric epithelial homeostasis (36). Of note, *NKX6-2* itself is predicted as a target for many of the transcription factors of the regulons with higher activity in *NKX6-2*+ areas, such as *FOXA3, NR2F6, GATA6*, and *KLF4* (**Table S15**).

Among the highest scored regulons negatively associated with *NKX6-2* expression, *ONECUT-2 and SPDEF* regulons were assigned higher activities in spots with no NKX6-2 expression. Both have been linked to PDAC pathogenesis via the non-cystic pancreatic intraepithelial neoplasia (PanIN) pathway and their overexpression is associated with adverse prognosis in invasive cancers (37,38). SPDEF is of particular interest because it has recently been implicated in the development of PDAC with classical differentiation and mucin production, two features that are also observed with elevated NKX6-2 in LG IPMNs (39). Nonetheless, the anti-correlation between NKX6-2 and SPDEF levels suggests that this transcription factor “switch” might be one of the prerequisites for progression of indolent LG lesions to higher grade lesions and cancer.

### Cross species validation of the gastric transcriptional program linked to Nkx6-2 expression

To evaluate *Nkx6-2* expression in a murine model of IPMN, we applied the Visium technology to analyze cystic lesions collected from two *Kras;Gnas* mice. Data was processed and 7 cluster assignments were identified (**Fig. S9A**). Based on gene expression and histologic assessment of the areas covered by the spots, clusters 0, 1, 3, and 6 were selected for further analyses of the epithelial compartments from pancreatic murine lesions (**Fig. S9B, Table S17**). Spots in these clusters were then grouped by *Nkx6-2* expression (*Nkx6-2* positive *versus Nkx6-2* negative) (**Fig. 6A**). Notably, higher expression of *Gkn1, Gkn2, Mucl3, Tff1, Tff2, and Muc5ac*, among others, which are markers of the NKX6-2-associated GIP signature identified in human IPMN samples, was observed in areas with *Nkx6-2* expression compared to those without detectable expression in the murine samples (**Fig. 6B, Table S18**). Gene set analyses of the differentially expressed transcripts between *Nkx6-2* positive *versus Nkx6-2* negative spots also showed enrichment of the human gastric pit-like cell profiles in areas with *Nkx6-2* expression (**Fig. S10A**). Expression of markers included in the GIP signature, such as *Tff1* and *Muc5ac*, have also been recently reported from single cell RNA sequencing analyses of murine pancreatic intraepithelial neoplasia (PanIN) cells in *Kras*^G12D^ induced acinar-to-ductal metaplasia models (37,38). On the other hand, *Reg1* and *Reg3b* levels were higher in areas with no detectable *Nkx6-2* expression. Genes encoding for pro-inflammatory proteins, such as *Cd74, C3*, and *C4b* were also found to be elevated in *Nkx6-2*-negative spots. Enrichment of these features have also been observed at later stages of progression in both acinar-derived (*Reg* genes) and ductal-derived cells (*Cd74, C3, C4b*) in *Kras*^G12D^ models compared to earlier time points (38). Interestingly, expression of *Cldn3, Igfbp7, Mmp7*, and *Ly6a*, which have been proposed as markers of a “pre-tumor” stage in murine lesions (37,38), were all detected at higher levels in areas of no *Nkx6-2* expression. Of note, many of these genes were also found to be negatively correlated to *NKX6-2* expression in the human ST IPMN dataset (**Fig. S10B**).

**Figure 6.**
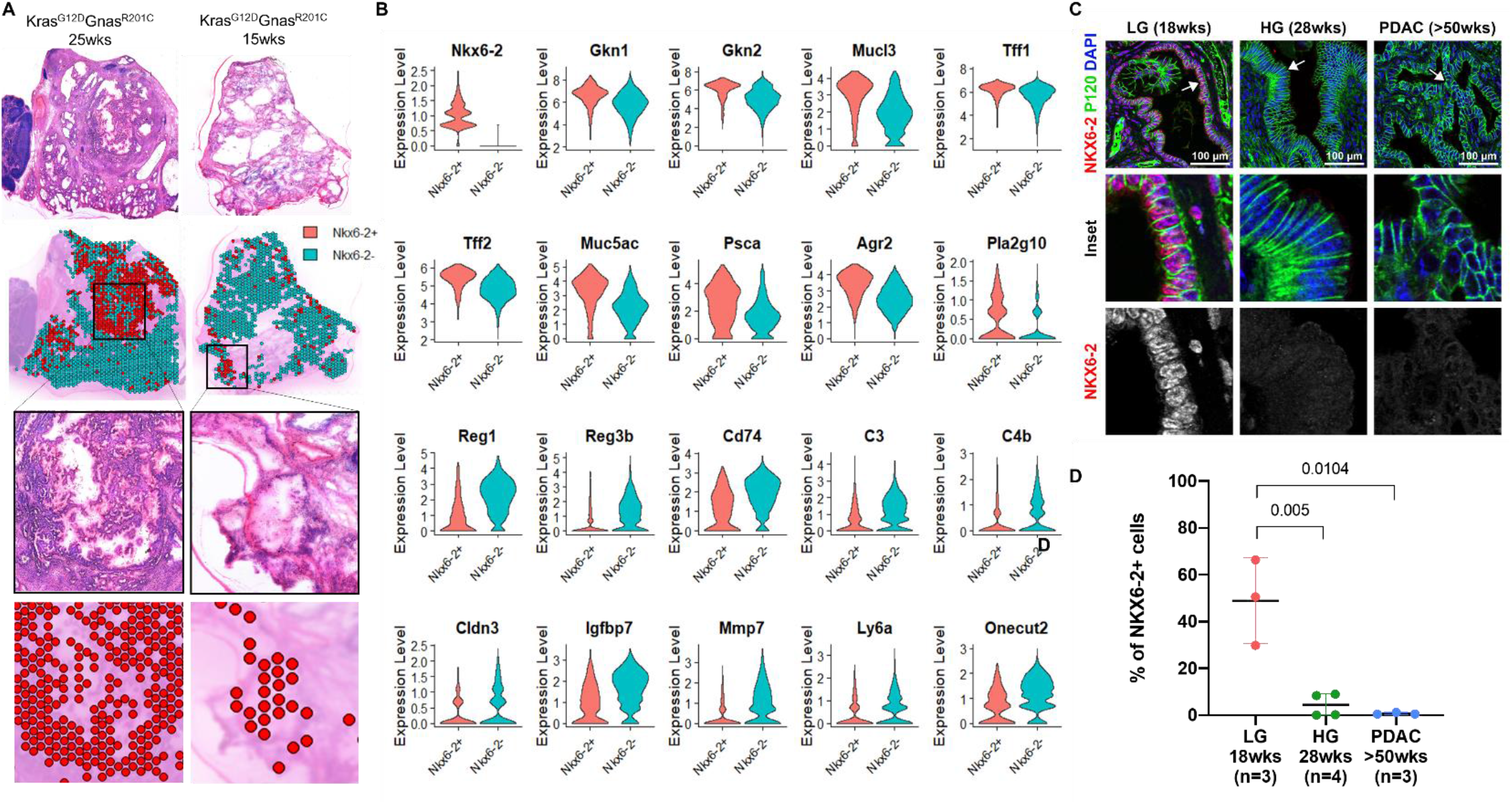
*Nkx6-2* expression in *a genetically engineered model* of IPMN. ST analyses of murine cystic lesions collected from Kras;Gnas mice. A) From top to bottom: H&E stained tissue samples. Spots corresponding to clusters of high epithelial contribution were selected and colored according to *Nkx6-2* expression, red = Nkx6-2+, blue = Nkx6-2 –. Zoomed-in images of areas presenting with high percentage of Nkx6-2 + spots. B) Expression of gene transcripts associated with *Nkx6-2* in the Kras;Gnas ST dataset shown in a violin plot. (C) Representative images of Kras;Gnas mouse tissues stained with NKX6-2 (red), P120 (green) and DAPI (blue). Arrows indicate enlarged area. (D) Quantification of the percentage of NKX6-2 positive cells in P120+DAPI+ epithelial cells. Three images per mouse captured under 40X magnification were used to count the number of cells and each dot shows the average percentage of cells in each mouse.

To evaluate a potential link between NKX6-2 expression and IPMN development and malignant progression, we stained for NKX6-2 with an epithelial cell marker (p120 catenin) and quantified the number of NKX6-2 positive epithelial cells in tissue samples collected from littermate *Kras;Gnas* mice after 18 weeks (n=3), 28 weeks (n=4), and 50+ weeks (n=3) on doxycycline diet (**Fig. 6C**). A significantly higher percentage of NKX6-2 positive cells was observed in the low-grade lesions from earlier stage mice compared to high-grade dysplasia and PDAC regions, supporting loss of NKX6-2 expression with tumor progression (**Fig. 6D**).

### Overexpression of NKX6-2 recapitulates the gastric transcriptional program and drives glandular differentiation in orthotopic models

To investigate gene expression and phenotypic changes that may be driven by *Nkx6-2*, we ectopically expressed *Nkx6-2* in a cell line isolated from the *Kras;Gnas* mice (**Fig. S11A**). The parental cell line (LGKC-5), derived from an established murine IPMN-associated cancer, as well as additional *Kras;Gnas*-derived cancer lines (LGKC-1 through -4), all lack discernible *Nkx6-2* expression (**Fig. S11B**). After lentivirus transduction, *Nkx6-2* expression was confirmed by qPCR and western blot analyses (**Fig. S11C-D**) and RNA-seq was performed in triplicate. A total of 7654 genes were found to be significantly differentially expressed by RNA-seq between the *Nkx6-2* overexpressing (Kras;Gnas^Nkx6-2+^) and control lines (**Table S19**). *Gkn2, Tff1, Mucl3, and Gkn1* were the four top ranked genes increased in *Nkx6-2* overexpressing cells. When comparing the differentially expressed transcripts between Kras;Gnas^Nkx6-2+^ and control cell lines to the transcriptomic profile associated with *Nkx6-2* expression in the murine ST dataset, 209 gene transcripts were found to exhibit overlapping trends, including higher levels of markers of the GIP signature (*Muc5ac, Tff1, Psca, Gkn2, Mucl3, Gkn1, Klf4*, and others) with *Nkx6-2* overexpression (**Table S20, Fig. 7A, Fig. S11E**). Remarkably, enrichment of gene sets corresponding to gastric isthmus and mature pit cells was also found by GSEA in the Kras;Gnas^Nkx6-2+^ cells, recapitulating our findings on the human ST datasets (**Fig. 7B**). A substantial overlap (n = 86 genes) was also observed between the GIP signature associated with *NKX6-2* expression in the human ST dataset, and the significantly altered transcripts driven by *Nkx6-2* overexpression in the *Kras;Gnas* cell lines (**Table S21, Fig. S11F-G**). Of note, ectopic expression of *Nkx6-2* in cell lines derived from the widely used *Kras;p53* (so-called “KPC”) mice, neither of which had any discernible *Nkx6-2* expression in the parental lines (**Fig. S11B, Fig. S12A-B**), similarly resulted in an enrichment of the gastric isthmus and pit cell gene sets by GSEA analyses (**Fig. S12C**). Several markers of the GIP signature (n=38) were found to be significantly enriched in the KPC^Nkx6-2+^ cell lines, including *Ctse, Vsig1, Lgals4*, and *Tspan8*. Overlapping trends in expression were also observed for 981 transcripts between the *Kras;Gnas* and *Kras;p53* cell lines upon *Nkx6-2* overexpression (**Table S22, S23**). Nonetheless, higher enrichment scores for the gastric gene sets were observed using the *Kras;Gnas* model, and some of the most characteristic genes of the GIP signature in IPMN, such as *Gkn1, Gkn2, Tff1*, among others, were not observed to be overexpressed in the KPC^Nkx6-2+^. Thus, the gastric transcriptional profile observed in the murine and human IPMN tissue samples was more closely recapitulated by ectopically expressing *Nkx6-2* in the *Kras;Gnas* compared to the *Kras;p53* model.

**Figure 7.**
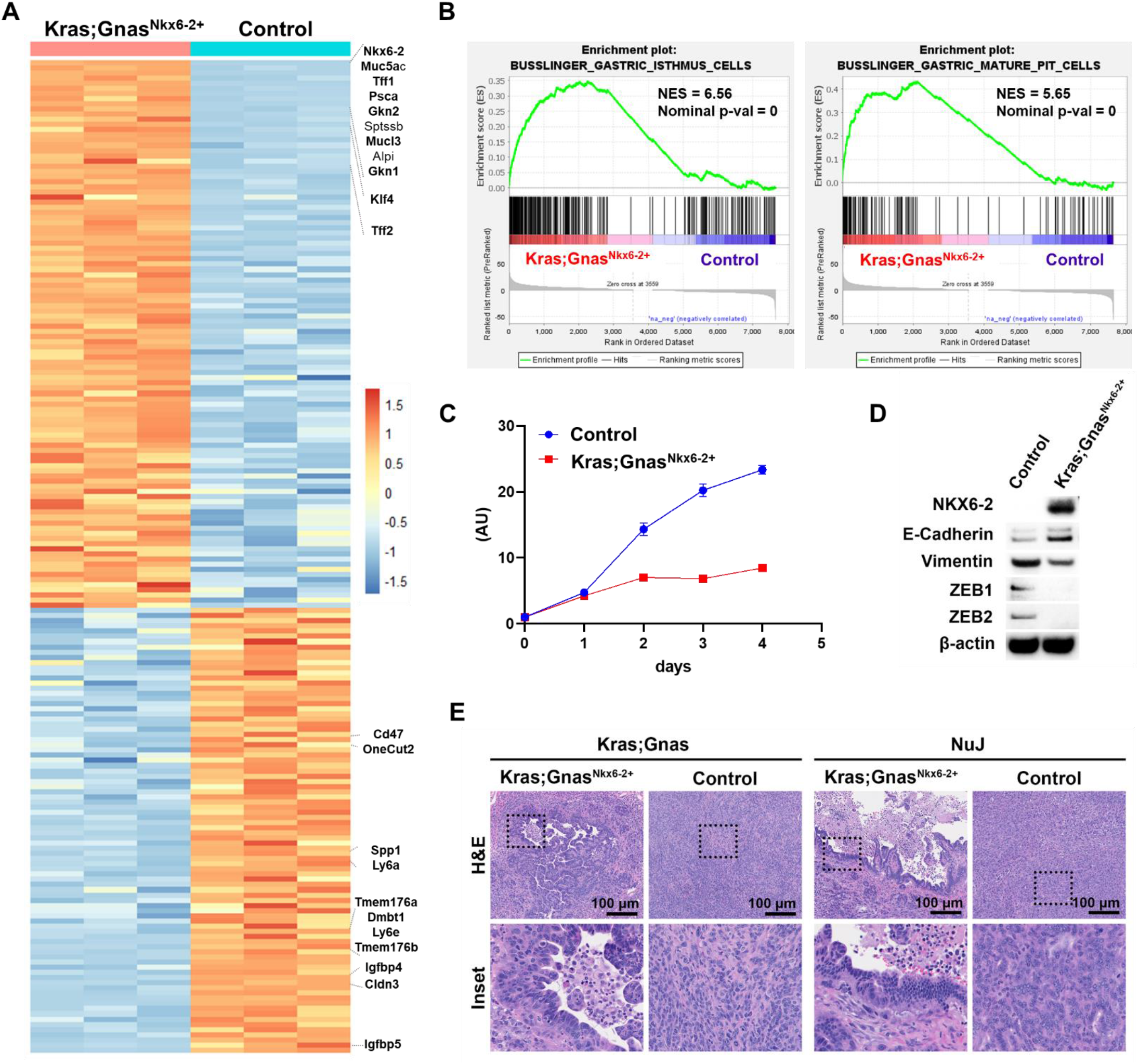
Enforced expression of Nkx6-2 in a cell line derived from Kras;Gnas mice. A) Heatmap showing normalized expression values of genes differentially expressed in the Kras;Gnas^Nkx6-2+^ versus control cell line that were also found to be differentially expressed in *Nkx6-2* + vs *Nkx6-2* – spots in the ST dataset from Kras;Gnas mice. Expression data is provided for three RNA-seq replicates. Note that genes with average raw counts < 10 from were not included in the heatmap. B) Enrichment of the gastric isthmus and pit cell gene sets in the *Nkx6-2* overexpressing cell line (Kras;Gnas^Nkx6-2+^). C) Cell proliferation assay using the Kras;Gnas^Nkx6-2+^ or control cells. * *P* < 0.05. D) Western blot of cell lysates obtained from the control and Kras;Gnas^Nkx6-2+^ cells, stained with NKX6-2, E-Cadherin, Vimentin, ZEB1, ZEB2 and β-actin antibodies. E) Representative H&E images of the tumors derived from the control or Kras;Gnas^Nkx6-2+^ cells at day 14 after orthotopic injection into pancreas of littermate (Kras;Gnas) or immunodeficient nude mice (NuJ). Images were captured under 10X or 40X (inset) magnification.

To evaluate how the induction of *Nkx6-2* affects cellular behavior, we performed a cell proliferation assay using the *Kras;Gnas* cell line overexpressing Nkx6-2 (*Kras;Gnas*^Nkx6-2+^). Decreased proliferation was observed in the *Kras;Gnas*^Nkx6-2+^ cells compared to control cells (**Fig. 7C**). Furthermore, western blots from the cell lysates showed higher expression of Vimentin, ZEB1, ZEB2 in the control cells, suggesting an overall increased EMT activity with NKX6-2 loss, as suggested by the transcriptomic data from the ST IPMN dataset (**Fig. 7D**). Additionally, we orthotopically implanted the *Kras;Gnas*^Nkx6-2+^ or parental (control) cells into the pancreas of littermate (Kras;Gnas) or immunodeficient athymic mice (NuJ). While both control and *Kras;Gnas*^Nkx6-2+^ cells formed orthotopic tumors within two weeks in both mouse strains on doxycycline diet, the histology of the tumors was strikingly different (**Fig. 7E**). The tumors established after injection of the *Kras;Gnas*^Nkx6-2+^ cells showed a significantly higher percentage of glandular structures (**Fig. S11H**), resembling IPMN-like papillary features, compared to the predominantly poorly differentiated tumors observed in the mice injected with control cells. This was a remarkable observation, suggesting that overexpression of a single transcription factor, NKX6-2, is capable of driving an IPMN-like glandular phenotype in an otherwise predominantly poorly differentiated histology. Overall, our cumulative cross species results support the role of *Nkx6-2* as a pivotal transcription factor in driving a gastric-like transcriptional profile in IPMNs, whose sustained expression might be a requirement for the observed indolent biological behavior of most gastric IPMN subtypes.

## Discussion

In the current study, ST analysis paired with single cell deconvolution provided novel insights into histologically annotated gene expression patterns in our cohort of patient derived archival IPMN samples. By using a region-specific approach of “epilesional”, “juxtalesional” and “perilesional” spots centered on proximity to the neoplastic epithelium, we were able to discern alterations in expression profiles that occur with IPMN progression within the neoplastic cells, as well as the surrounding milieu. While several microenvironment-specific alterations in immune cell composition were observed on ST, the validation studies performed with the COMET cyclic IF platform on an independent cohort of archival IPMNs showed a similar trend in the reduction of plasma cell infiltration with progression. This suggests that humoral immunity may be an important mediator of immune surveillance in early stages of multistep IPMN progression, and this function wanes with subsequent progression to HG dysplasia and cancer. As recently shown by Mirlekar and colleagues, the balance between plasma cells and immune suppressive regulatory B cells governs immune surveillance in established PDAC, with our analysis supporting a role for the former even in non-invasive disease (40). Mast cells were also found to be an abundant component of the IPMN microenvironment (varying between 5-10% of the immune cells), albeit the COMET platform did not show a significant attenuation in mast cell numbers with IPMN progression. Mast cells have been implicated in the development and growth of “conventional” PDAC, but their role in cystic precursor lesions, especially in early disease, remains to be explored (41).

In this ST analysis of a panel of IPMNs, we identified the homeobox transcription factor *NKX6-2* to be a pivotal driver of a so-called “gastric isthmus-pit” (GIP) signature in LG IPMNs of the gastric subtype, the most common subtype of mucinous cystic lesion of the pancreas. NKX6-2 protein levels were also quantified by COMET cyclic IF in an independent cohort of IPMN archival samples, showing increased levels of nuclear NKX6-2 in the epithelium of LG IPMN compared to HG IPMN, together with co-expression of GIP markers MUC5AC and TFF1. Leveraging the spot-level and full-transcriptome information provided by Visium, we investigated the transcriptomic profiles associated with *NKX6-2* expression, which resulted in the identification of a highly enriched signature resembling that of gastric isthmus and pit cells of the stomach, which we termed as GIP signature. Even though gastric IPMN is morphologically similar to gastric epithelium, the remarkable similarity between the *NKX6-2* expressing LG gastric IPMN and the gastric stomach isthmus-pit cell transcriptome was striking. The *NKX6-2* associated GIP signature was also observed in an independent RNA sequencing cohort of archival IPMN samples conducted as part of the NCI Cancer Moonshot Precancer Atlas Pilot project (14). Moreover, *Nkx6-2* expression was also observed in IPMN lesions collected from a genetically engineered IPMN model (13), with enrichment of the GIP signature also detected in spots of high *Nkx6-2* expression. Loss of Nkx6-2 was also associated with the progression of IPMN lesions in the *Kras;Gnas* mice as determined by IF, with higher percentage of NKX6-2 positive cells in low-grade compared to lesions of high-grade dysplasia and cancer in those samples.

To investigate the role of *NKX6-2* in driving the observed gastric IPMN phenotype, we overexpressed *Nkx6-2* in an IPMN-derived cancer cell line generated from the *Kras;Gnas* mouse model. Remarkably, this single alteration alone was able to recapitulate the GIP signature observed with *NKX6-2* in both human and murine IPMN tissue datasets and presented a more indolent phenotype with a decrease in cellular proliferation and EMT activity. Further, the histology of the *Nkx6-2* overexpressing orthotopic tumors was substantially distinct from the parental cells, with an obvious increase in the percentage of glandular features resembling that of IPMN in the Kras;Gnas^Nkx6-2+^ tumors. Overexpression of *Nkx6-2* in *Kras;p53* cell lines also resulted in enrichment of the GIP signature, albeit with less overlap to the transcriptomic profiles associated to NKX6-2 expression in murine and human IPMN compared to the Kras;Gnas^Nkx6-2+^ cells. This was not an unexpected finding, given that the *Kras;Gnas* model closely mimics IPMN development, while the *Kras;p53* model leads to PanIN and PDAC lesions. Overall, we hypothesize that a conserved transcriptional program driven by *NKX6-2* exists independently of the *TP53* or *GNAS* mutant status, but that the co-existing genomic alternations might be impacting the extent of downstream effector pathways that are modulated. Ongoing experiments modulating *Nkx6-2* expression in autochthonous models of IPMNs will help elucidate the role and mechanism of this transcription factor in upregulating the GIP signature and driving glandular differentiation.

Although *NKX6-2* expression is consistently higher in LG IPMNs *versus* HG IPMNs and invasive cancer, including in cases with synchronous lesions of distinct histological grades, we did observe heterogeneity of *NKX6-2* expression within LG IPMNs. Previous single cell RNA sequencing analyses performed by our group have also identified intra-lesion heterogeneity in IPMN, with a minor subpopulation of epithelial cells in LG IPMN demonstrating transcriptomic overlap with HG IPMNs or even cancer (42). This leads us to hypothesize that while elevated *NKX6-2* expression is an accompaniment of the earliest steps of IPMN development, loss of NKX6-2 within the LG epithelium might be a harbinger of progression to HG IPMN or cancer. The assessment of differentially expressed genes and transcriptional regulatory networks between *NKX6-2* positive and negative spots within LG IPMNs were quite revealing in this regard. For example, epilesional spots with minimal *NKX6-2* expression in LG IPMNs showed higher levels of PDAC-associated markers, such as *CD74, CXCL2, LY6E*, and *CLDN4*, as well as *KRT17*, a marker of the aggressive basal subtype of PDAC, pointing to a transcriptional reprogramming away from the indolent NKX6-2-expressing epithelium (18,27,28,31). On the other hand, enrichment of the classical-like signature was observed in spots with high NKX6-2 expression, with higher levels of GATA6 staining quantified in NKX6-2-expressing nuclei by cyclic IF (15,30). Similarly, SCENIC analysis for aberrant regulons found that even in LG IPMNs, loss of *NKX6-2* leads to higher predicted activities of ONECUT-2 and SPDEF regulatory networks, both markers of PDAC progression and aggressive biology (37-39). The cumulative assessment from this data leads us to believe that *NKX6-2* serves as an “epithelial rheostat” in IPMN pathogenesis, and its loss acts as a transcription factor switch, driving an otherwise indolent mucin producing epithelium on a path towards invasive neoplasia.

Our study has certain limitations. First, it is important to note that all IPMN cases included in this study were surgically resected and thus, by definition, the natural history of the lesions was disrupted. Therefore, determining the impact of longitudinal alterations in *NKX6-2* expression within the cyst epithelium on clinical outcome was not feasible. In future studies, such effect could be evaluated by potentially monitoring methylation levels of *NKX6-2* DNA in cystic fluid of LG IPMNs that are undergoing surveillance. Loss of *NKX6-2* expression has been previously attributed to DNA methylation events, and detection of methylated *NKX6-2* DNA in biofluids has been proposed as an early detection biomarker for various cancer types, such as in bladder and renal carcinomas (43,44). Second, the Visium technology used in this study did not permit the capture of transcripts at single cell resolution, but the application of deconvolution approaches as well as targeted selection of tissue areas allowed us to capitalize on the available histological information to circumvent this challenge. Moreover, the use of the innovative cyclic IF COMET technology enabled the quantification and spatial localization of selected epilesional and juxtalesional markers at the protein level. Finally, an important unanswered question in our study remains the upstream regulators of *NKX6-2* expression in gastric IPMNs.

Given the heterogeneity of IPMN lesions in the pancreas, efforts directed at accurately defining the molecular events dictating biological potential and propensity for progression to cancer are paramount for improving their clinical management. The establishment of molecular markers associated with an indolent biology will prevent the overtreatment of otherwise benign lesions, while encouraging increased surveillance and treatment of higher risk lesions. We posit that NKX6-2 is not only a driver of the unique gastric transcriptional profile and histological phenotype that is observed in early IPMNs, but its sustained expression in the IPMN epithelium portends an indolent biological potential in this cystic precursor lesion.

## Materials and Methods

### Patient and murine sample collection for ST analyses

Archival snap frozen (**Table S1**) tissue samples from 13 IPMN resections at either MDACC or Indiana University School of Medicine were sectioned, stained with hematoxylin & eosin (H&E) and evaluated by a pancreas pathologist (A.M) to identify areas containing IPMN lesions and cancer. The analyzed samples were comprised of 7 LG IPMN (all gastric), 3 HG IPMN, and 3 PDAC arising in the backdrop of IPMN. All biospecimens were collected under IRB approved protocols. At MDACC, tissue samples were collected under the Pancreas Tissue Bank, LAB00-396 and subsequently analyzed under a use protocol PA15-0014. For IU, the biospecimens were collected under protocol 1011003217 (0209-66). This study was conducted in accordance with Good Clinical Practices concerning medical research in humans per the Declaration of Helsinki. Samples were cryo-sectioned onto Visium Spatial Gene Expression slides for ST analysis. For the murine samples, pancreata were collected from p48-Cre; LSL-KrasG12D; Rosa26R-LSL-rtTA-TetO-GnasR201C mice (*Kras*;*Gnas* mice) at 15 and 25 weeks on doxycycline diet. Samples were flash frozen in liquid nitrogen, cryo-sectioned and H&E stained. After sectioning onto the Visium slides, sample processing was performed as described in the Visium 10x Genomics protocol for fresh-frozen tissue samples (15 minutes were used for permealization time). After library preparation, samples were pooled and sequenced using a NextSeq 550 (Ilumina, CA, USA). The sequencing outputs for all Visium datasets were preprocessed using SpaceRanger and mapped to either the latest human or mouse reference genome and the filtered matrices were converted to Seurat objects in R and used for downstream analyses.

### ST data analysis

The single-cell data integration method to match and compare (scMC) (16) pipeline was used for initial processing and clustering of the human IPMN ST dataset. To infer single cell composition from the ST dataset, we applied the robust cell type deconvolution (RCTD) algorithm (17) using full-mode and a previously collected and annotated single-cell RNA sequencing pancreatic cancer dataset as reference (45), which provided cell type proportions for each of the spots. For histologically directed analysis, the Loupe Browser software was used to select spots covering specific areas in the tissue samples. ST data for the two *Kras;Gnas* samples was SCT normalized and merged. Differential expression analyses of ST data were performed using the FindMarkers and FindAllMarkers functions in Seurat, filtering based on Log2FC, minimum percentage of spots expressing that gene, and adjusted p-value (log2FC > 0.25, min.pct > 0.25, p-val_adj < 0.05). GSEA analyses were conducted using the GSEA 4.2.3 software and the ranked features from differential expression. To evaluate changes in cell composition in the three selected compartments, log2FC and p-values of the per-spot cell-type proportions between LG and HG+PDAC samples were calculated. For regulon analyses, we applied the Single-cell rEgulatory Network Inference and Clustering (SCENIC) (33) method to evaluate regulatory networks that might be associated with NKX6-2 expression. *GENIE3* was first used to identify genes co-expressed with transcription factors based on the transcriptional profiles obtained from epilesional areas in the LG IPMN samples (32). These co-expression modules were then evaluated using *RcisTarget* based on cis-regulatory motif analyses to identify regulons. The activity of each regulon was weighted per each spot and compared between the areas with and without detected *NKX6-2* expression. To evaluate differences in regulon activity associated to *NKX6-2* expression, regulons were scored based on the fold change in activity between the two groups and average activity value per group.

### COMET cyclic IF protein profiling

We performed immunofluorescence staining of 19 protein markers, which were selected based on transcriptomic changes observed from the ST dataset (**Table S24**), on an independent cohort of 26 archival FFPE IPMN tissue samples (**Table S9**) using the COMET platform (Lunaphore, Switzerland) (*see supporting methods*). Prior to imaging and staining, antigen retrieval was performed using EZ antigen retrieval buffer at 107°C for 15 minutes in an EZ-Retriever Microwave System (BioGenex, CA, USA). Images were imported into Visiopharm software, where IPMN areas were selected, and nuclei and cell segmentation were performed for quantification (*see supporting methods*).

### Overexpression of Nkx6-2 in a murine Kras;Gnas and KPC cell lines

We used a cell line derived from a murine IPMN-associated PDAC in a 12-month-old male *Kras;Gnas* mouse (13) (p48-*Cre*;LSL-Kras^G12D^;Rosa26R-LSL-rtTA-TetO-Gnas^R201C^, on 0.0060% doxycycline diet) and two conventional KPC (K-ras^LSL.G12D/+^; and Trp53^R172H/+^; Pdx-1-Cre) cell lines. A Nkx6-2 vector: pLV[Exp]-Puro-CMV>mNkx6-2[NM_183248.3](ns)/3xFLAG/Myc:T2A:EGFP or a control vector (pLV[Exp]-EGFP:T2A:Puro-EF1A>mCherry) (VectorBuilder Inc, IL, USA) were transduced in the cell lines, followed by puromycin (Thermo Fisher Scientific, Inc., MA, USA) selection. After the construction of Nkx6-2 overexpressing cell line, we performed WST-8 assay (Nacalai Tesque, Kyoto, Japan) to analyze cell proliferation according to the manufacturer’s protocol.

### Orthotopic injection

Immunodeficient nude mice (NuJ) purchased from The Jackson Laboratory (Bar Harbor, ME, US) or Kras;Gnas littermate mice from which the cell line was derived were used for orthotopic injection. The detailed procedure was described previously (46). Briefly, 1 × 10^6^ cells (NKX6-2 overexpressing or empty vector-containing control) resuspended in gel matrix were injected orthotopically into the body of the pancreas under general anesthesia. The experiment was conducted in compliance with the Institutional Animal Care and Use Committee (IACUC) guidelines at MD Anderson Cancer Center (MDACC).

## Acknowledgements

The authors would like to thank Drs. Vittorio Branchi, Maria Monberg, Sonja Woermann, Cara Haymaker, and Mr. Bret M. Stephens for helpful discussions. The authors acknowledge Nathaniel Yee and Dorsay Sadeghian for assistance.

## Data Availability

The Visium and bulk RNA sequencing datasets are uploaded to the NCBI Gene Expression Omnibus (GEO) database.

