## Supplementary figures and methods for "Spatial Transcriptomics of Intraductal Papillary Mucinous Neoplasms of The Pancreas Identifies NKX6-2 as a Driver of Gastric Differentiation and Indolent Biological Potential"

**Supporting Information**

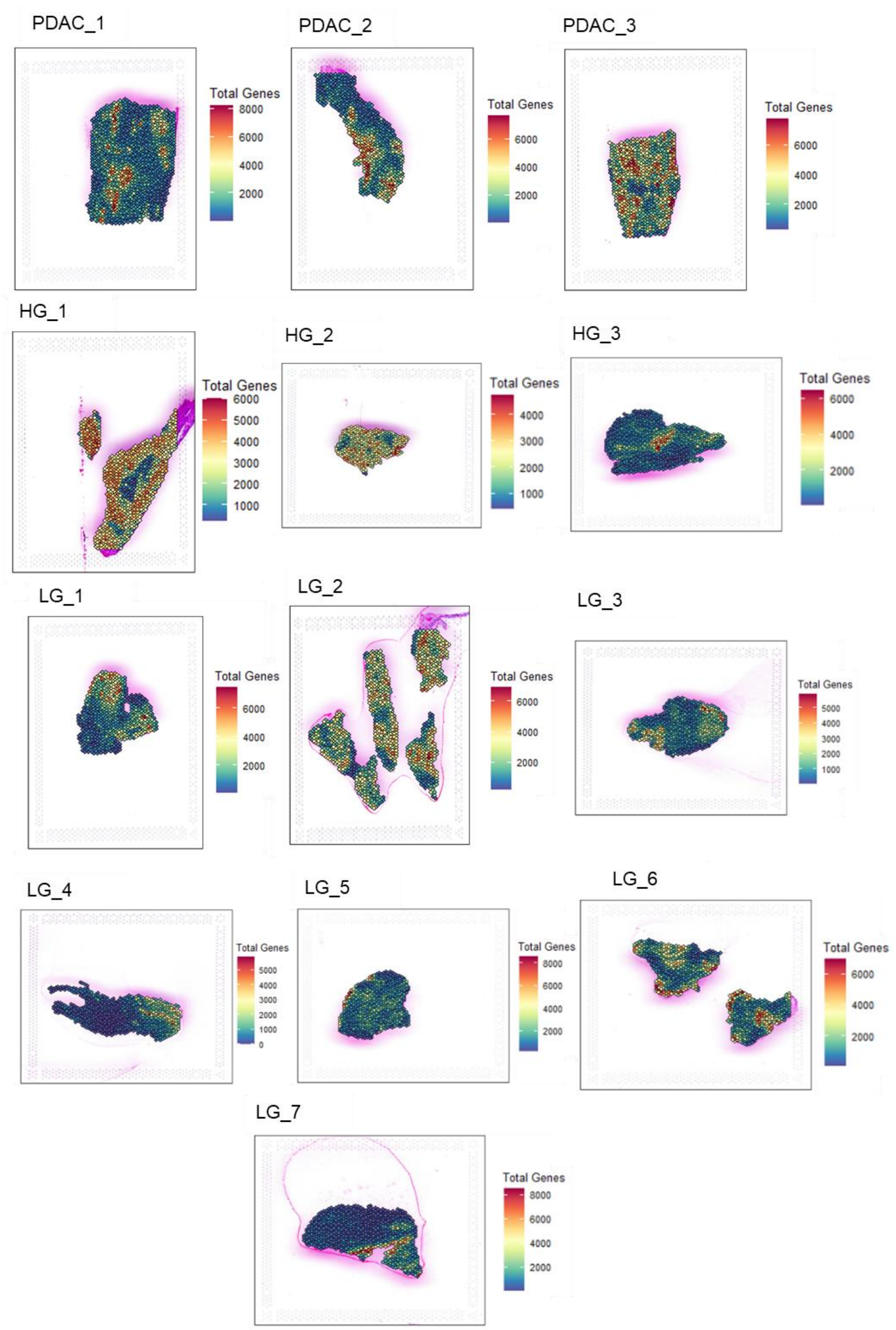


**Figure S1**. Spatial maps showing number of genes detected per spot in the 13 samples used for spatial transcriptomics analyses. Data was collected from a total of 12685 spots with up to 8000 genes detected per each spot.

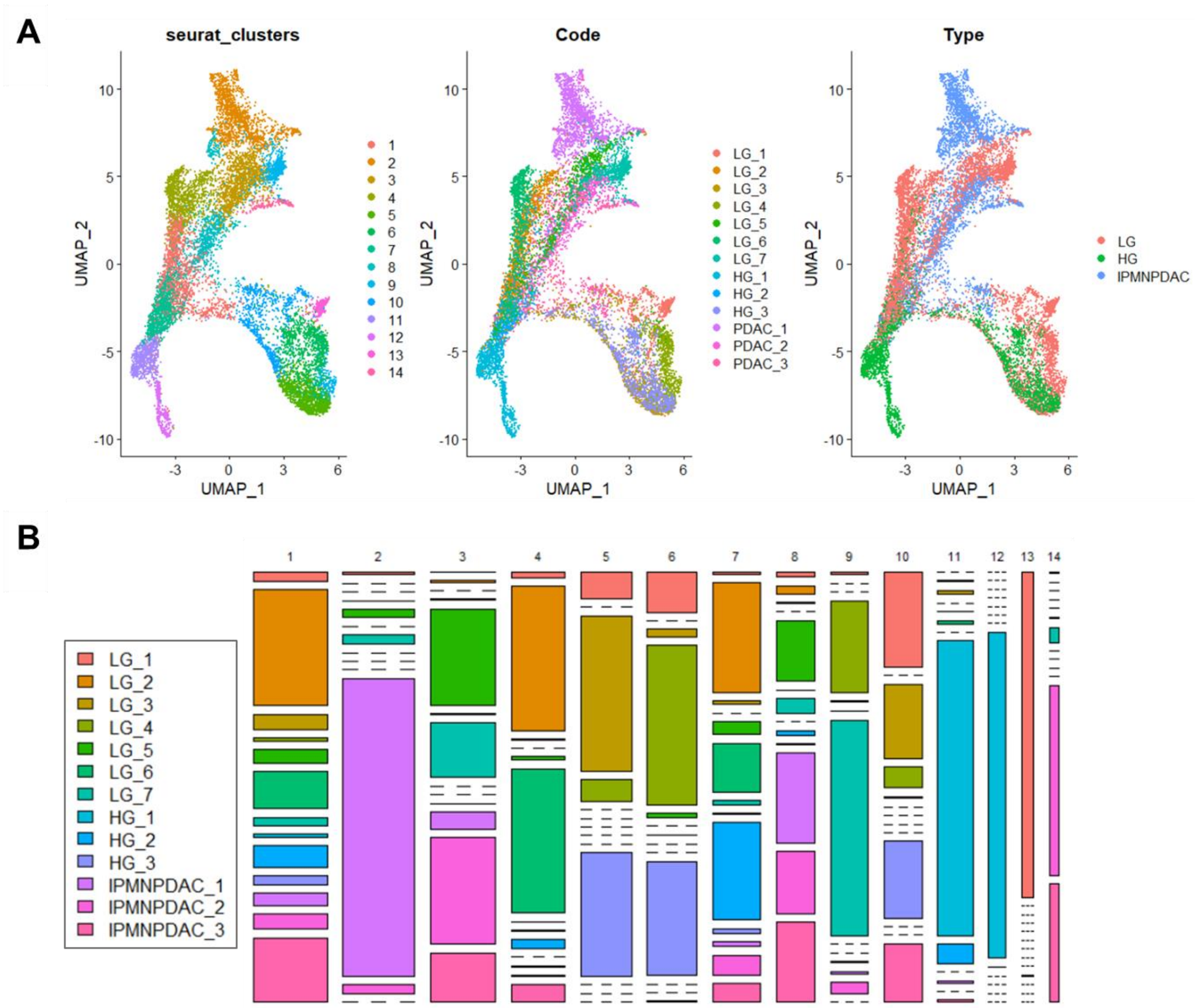


**Figure S2**. A) Uniform Manifold Approximation and Projection (UMAP) of all spots collected from the 13 samples by ST. Spots are colored according to transcriptomic cluster (seurat_clusters), sample name, and sample type – LG: low-grade IPMN, HG: high-grade IPMN, IPMNPDAC: high-grade IPMN with co-occurring PDAC. B) Sample proportions for each of the 14 transcriptomic clusters defined by ST and scMC analyses.

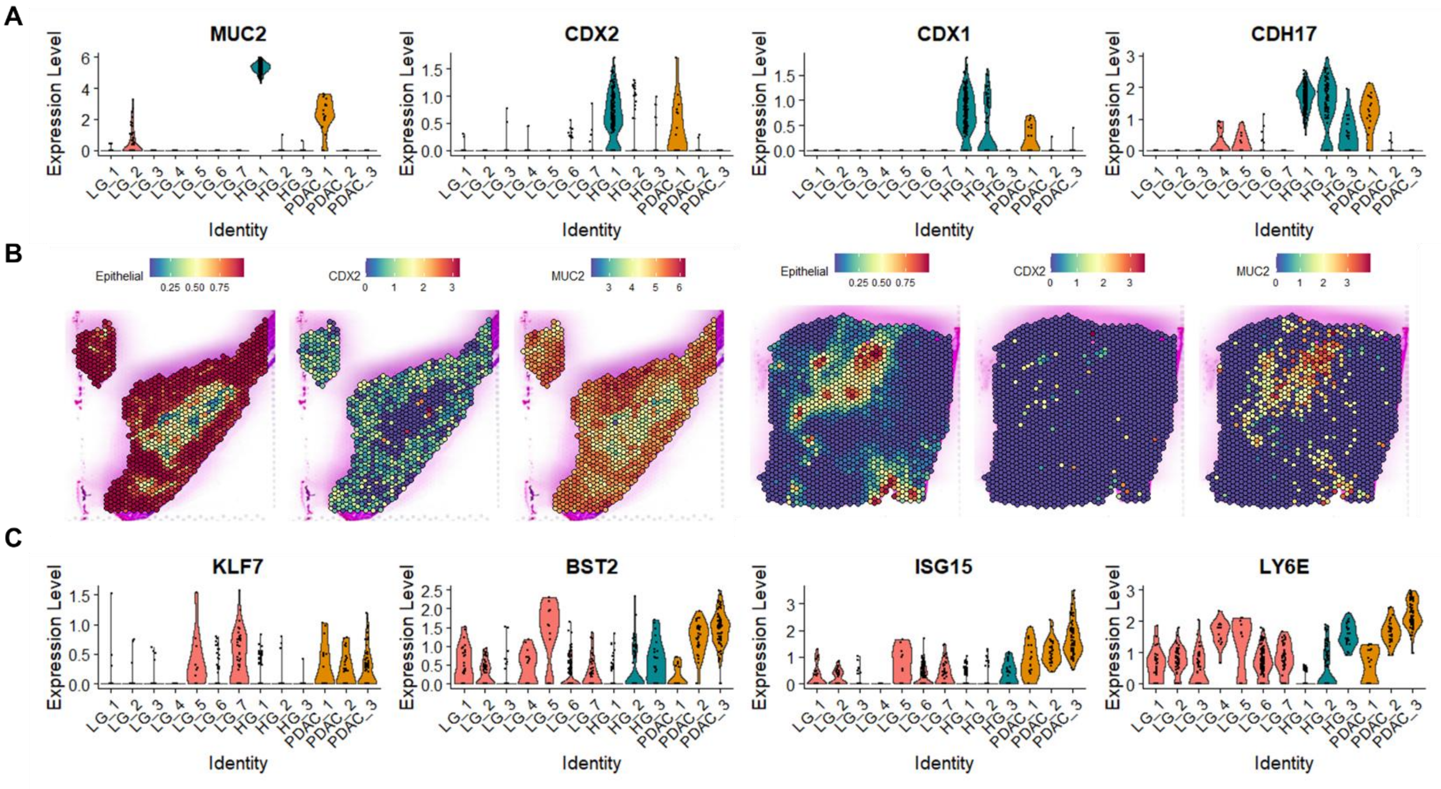


**Figure S3**. A) Violin plot showing expression of CDX2 and MUC2 in the epilesional compartments of IPMN and PDAC samples. B) Spatial maps showing the localization and proportions of epithelial cells per spot, as well as expression and localization of the CDX2 and MUC2 transcripts in sample HG_1 (left, HG intestinal case), and PDAC_1 (right, colloid carcinoma). C) Violin plots showing expression of transcripts with increased expression in PDAC compared to LG and HG IPMN epilesional spots.

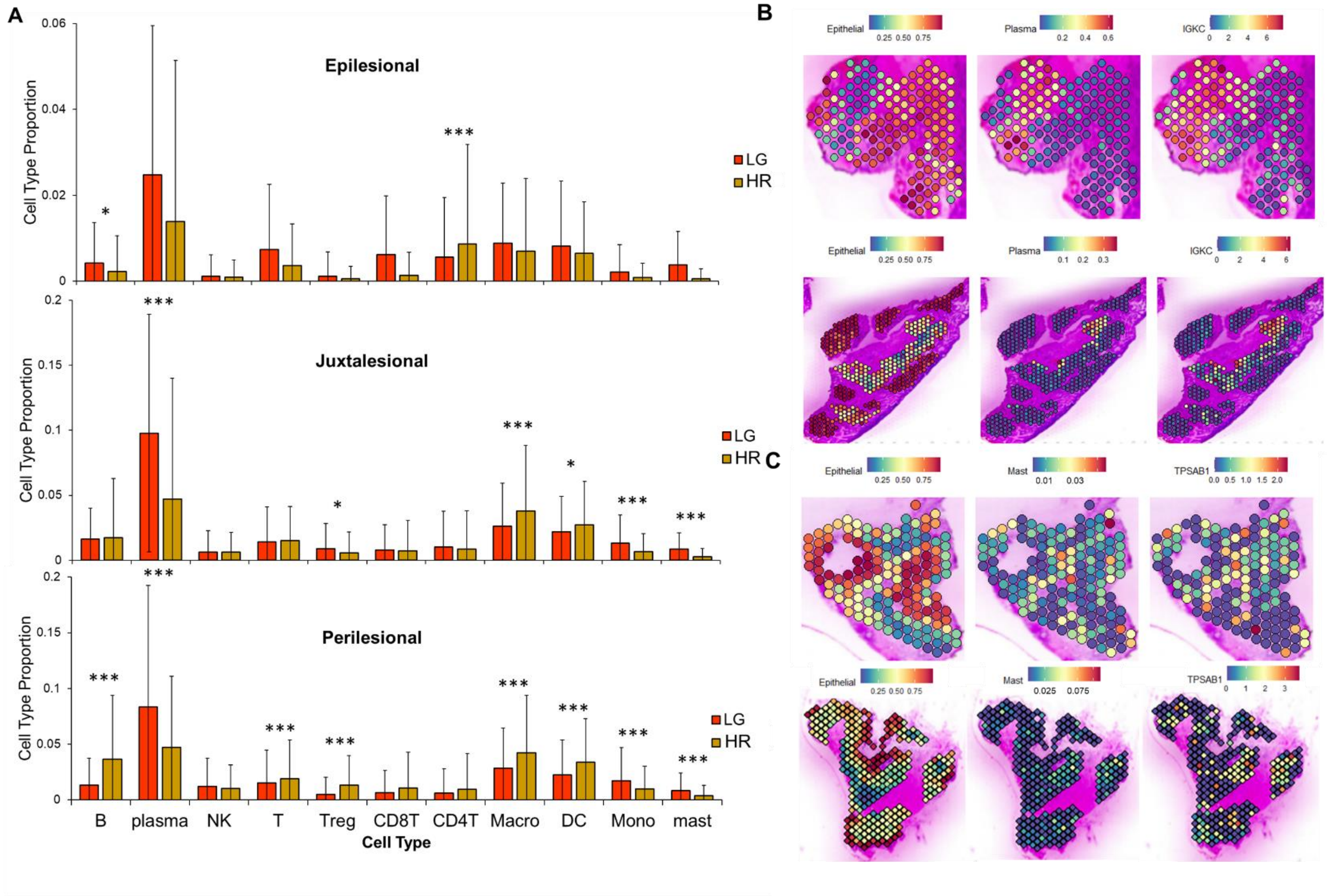


**Figure S4**. **Cell type proportions in LG and HR IPMN compartments determined by RCTD deconvolution**. A) Bar plot showing average cell type proportions in the epilesional, juxtalesional, and perilesional compartments in LG and HR IPMN samples. * $p < 0.05$, ** $p < 0.01$, *** $p < 0.001$. Expression and co-localization of predicted cell proportions by RCTD with markers of the selected cell type for (B) plasma and (C) mast cells.

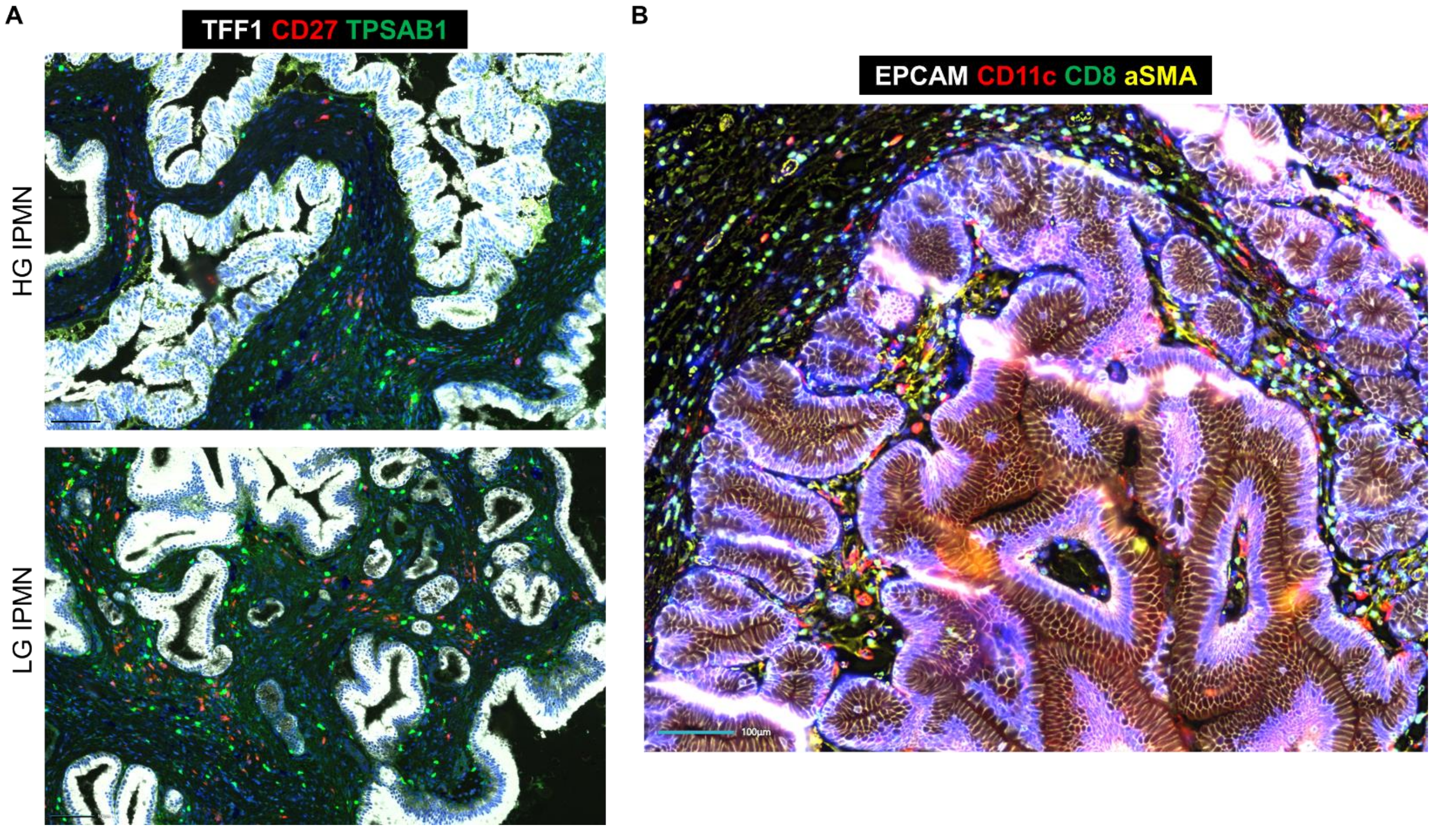


**Figure S5**. **COMET cyclic IF imaging of IPMN**. A) Plasma (CD27) and mast (TPSAB1) cell staining in areas of synchronous HG (top) and LG (bottom) IPMN areas from the same tissue section. B) COMET imaging of a LG IPMN showing additional markers evaluated for TME profiling in IPMN.

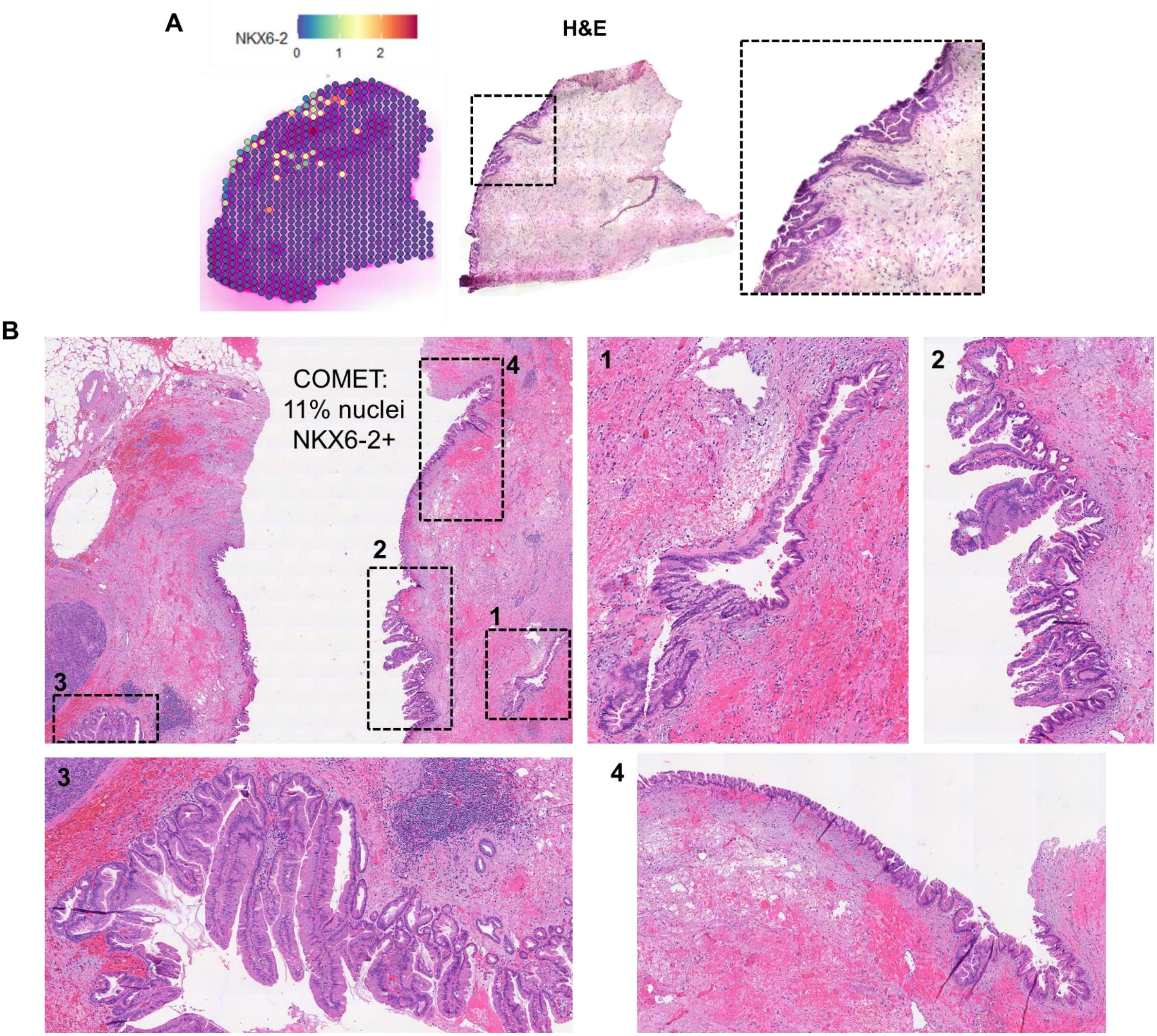


**Figure S6**. **NKX6-2 levels determined by ST and COMET analyses conducted on tissue samples from the same patient**. A) Spatial map showing levels of *NKX6-2* transcript in sample LG_5 of the ST dataset. An H&E of the same tissue is also provided showing in the zoomed-in image a LG gastric IPMN in a potential transitional state to pancreatobiliary IPMN. B) H&E images showing the areas analyzed in the LG_1 case by COMET imaging. COMET analyses and subsequent quantification resulted in the identification of 11% of NKX6-2 positive nuclei in the IPMN areas selected. Zoomed-in H&E images of areas quantified for NKX6-2, which also showed potentially transitionary states between gastric and pancreatobiliary IPMN.

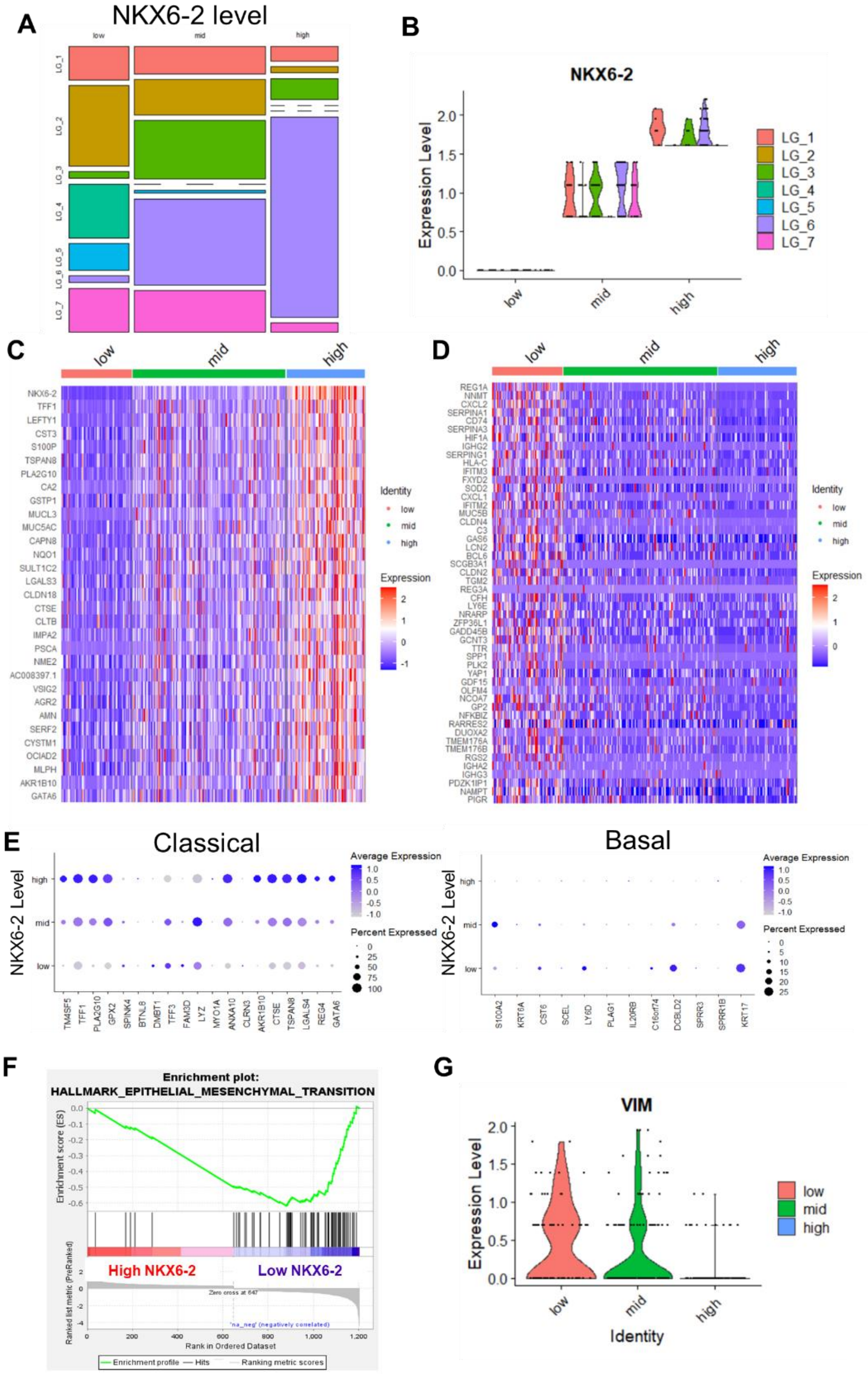


**Figure S7**. ***NKX6-2* expression in LG IPMN samples and its associated signature**. A) Proportion of spots per each of the *NKX6-2* level groups: low, mid, and high, split by sample name for the LG IPMN tissues. B) Split of LG epilesional spots according to *NKX6-2* expression (low: expression level = 0, mid: expression level > 0.5 & < 1.5, high; expression level > 1.5). Heatmaps showing normalized expression of gene transcripts associated with increasing (C) or decreasing (D) *NKX6-2* levels. The genes included in B were obtained from selecting features that were differentially expressed between all comparisons (high vs low, high vs mid, mid vs low). Genes included in C were obtained from differential expression analyses between spots with minimal *NKX6-2* expression (low) versus spots with mid and high *NKX6-2* expression. E) Expression of prototypal genes from the classical (left) and basal (right) PDAC subtypes in the *NKX6-2* low, mid, and high groups, visualized in a dot plot. F) Enrichment plot of the epithelial to mesenchymal transition gene set obtained through ranked GSEA analyses using the differentially expressed genes between low and high (high+mid) *NKX6-2* spots. G) Violin plot showing *VIM* levels in LG IPMN epilesional areas split by *NKX6-2* levels.

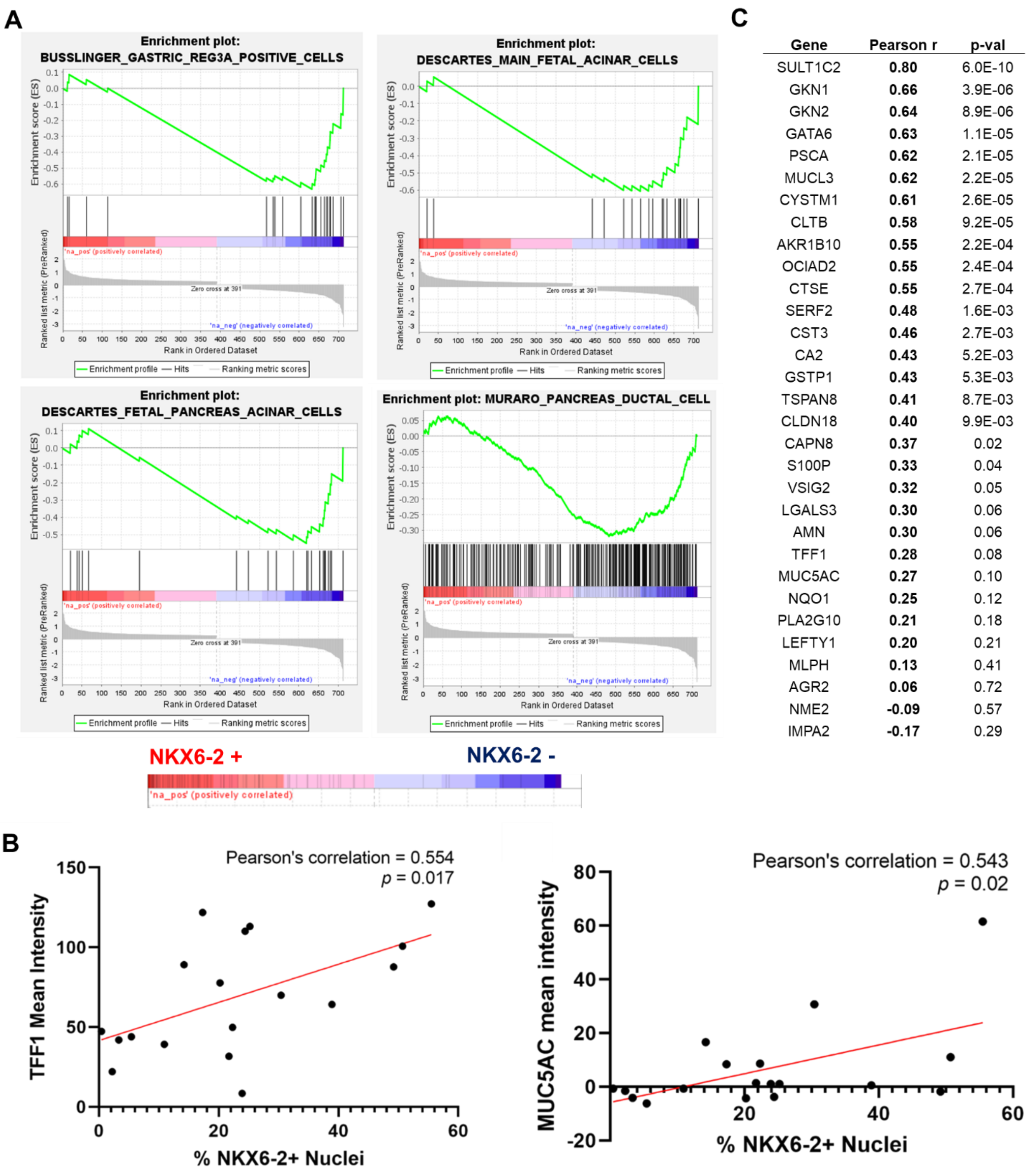


| Gene | Pearson r | p-val |
|---|---|---|
| SULT1C2 | **0.80** | 6.0E-10 |
| GKN1 | **0.66** | 3.9E-06 |
| GKN2 | **0.64** | 8.9E-06 |
| GATA6 | **0.63** | 1.1E-05 |
| PSCA | **0.62** | 2.1E-05 |
| MUCL3 | **0.62** | 2.2E-05 |
| CYSTM1 | **0.61** | 2.6E-05 |
| CLTB | **0.58** | 9.2E-05 |
| AKR1B10 | **0.55** | 2.2E-04 |
| OCIAD2 | **0.55** | 2.4E-04 |
| CTSE | **0.55** | 2.7E-04 |
| SERF2 | **0.48** | 1.6E-03 |
| CST3 | **0.46** | 2.7E-03 |
| CA2 | **0.43** | 5.2E-03 |
| GSTP1 | **0.43** | 5.3E-03 |
| TSPAN8 | **0.41** | 8.7E-03 |
| CLDN18 | **0.40** | 9.9E-03 |
| CAPN8 | **0.37** | 0.02 |
| S100P | **0.33** | 0.04 |
| VSIG2 | **0.32** | 0.05 |
| LGALS3 | **0.30** | 0.06 |
| AMN | **0.30** | 0.06 |
| TFF1 | **0.28** | 0.08 |
| MUC5AC | **0.27** | 0.10 |
| NQO1 | **0.25** | 0.12 |
| PLA2G10 | **0.21** | 0.18 |
| LEFTY1 | **0.20** | 0.21 |
| MLPH | **0.13** | 0.41 |
| AGR2 | **0.06** | 0.72 |
| NME2 | **-0.09** | 0.57 |
| IMPA2 | **-0.17** | 0.29 |

**Figure S8**. A) Enrichment plots for gastric REG3A positive cells, main fetal acinar cells, pancreas acinar cells, and ductal cells obtained via GSEA analyses. B) Correlation between % NKX6-2 positive nuclei and TFF1 or MUC5AC expression in the COMET images. The red line represents the linear regression line. C) Correlation analyses conducted using the RNA sequencing PCAPP dataset of archival IPMN tissue samples. Pearson correlation coefficients and p-values were calculated between NKX6-2 and representative genes of the associated GIP signature shown in Fig. 5C.

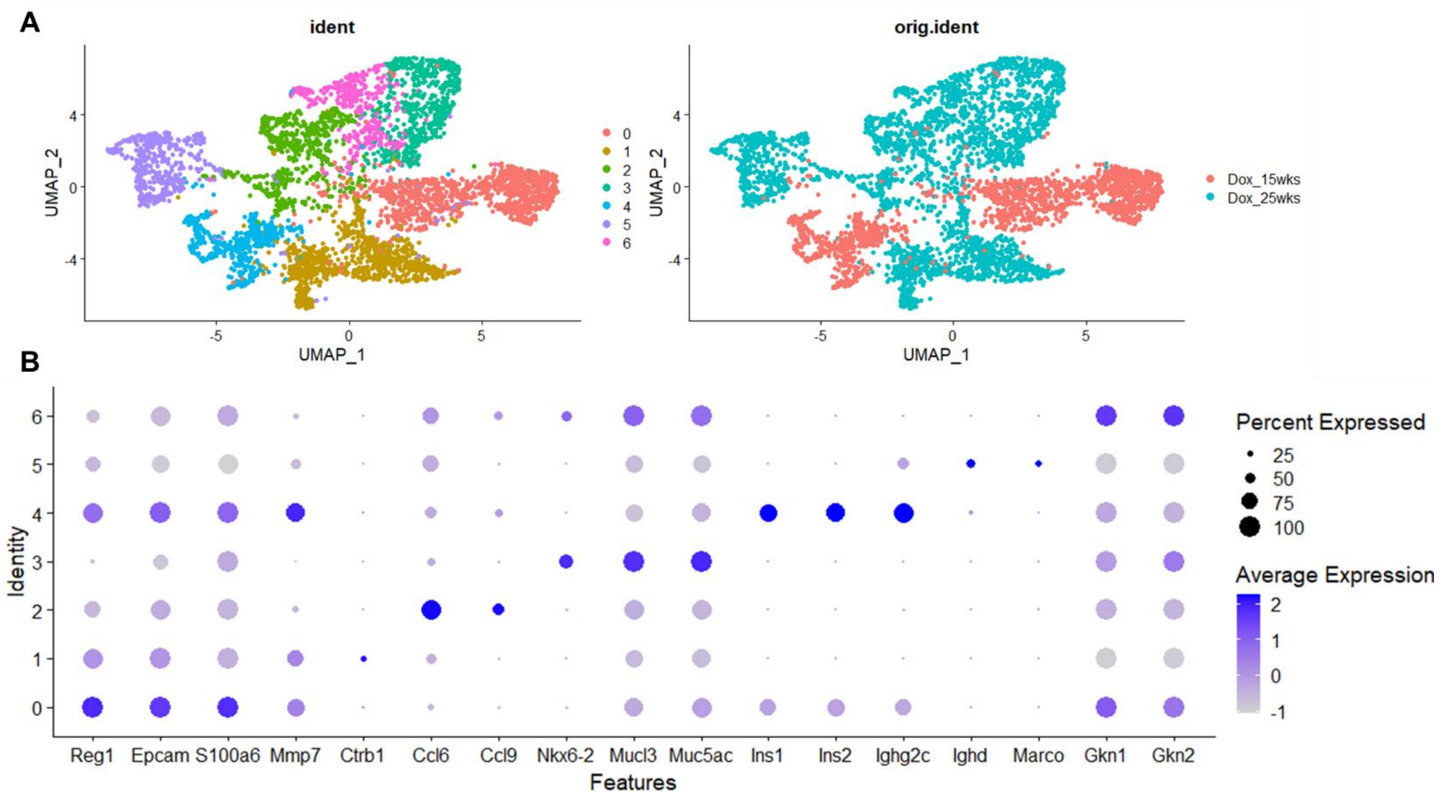


**Figure S9**. **Spatial transcriptomic analyses of a murine model of IPMN**. A) Uniform Manifold Approximation and Projection (UMAP) of the spots obtained from ST analyses of tissue samples from Kras;Gnas mice. Spots are colored according to cluster assignments (left) and sample of origin (right). B) Dot plot of top differentially expressed markers between each of the 7 clusters.

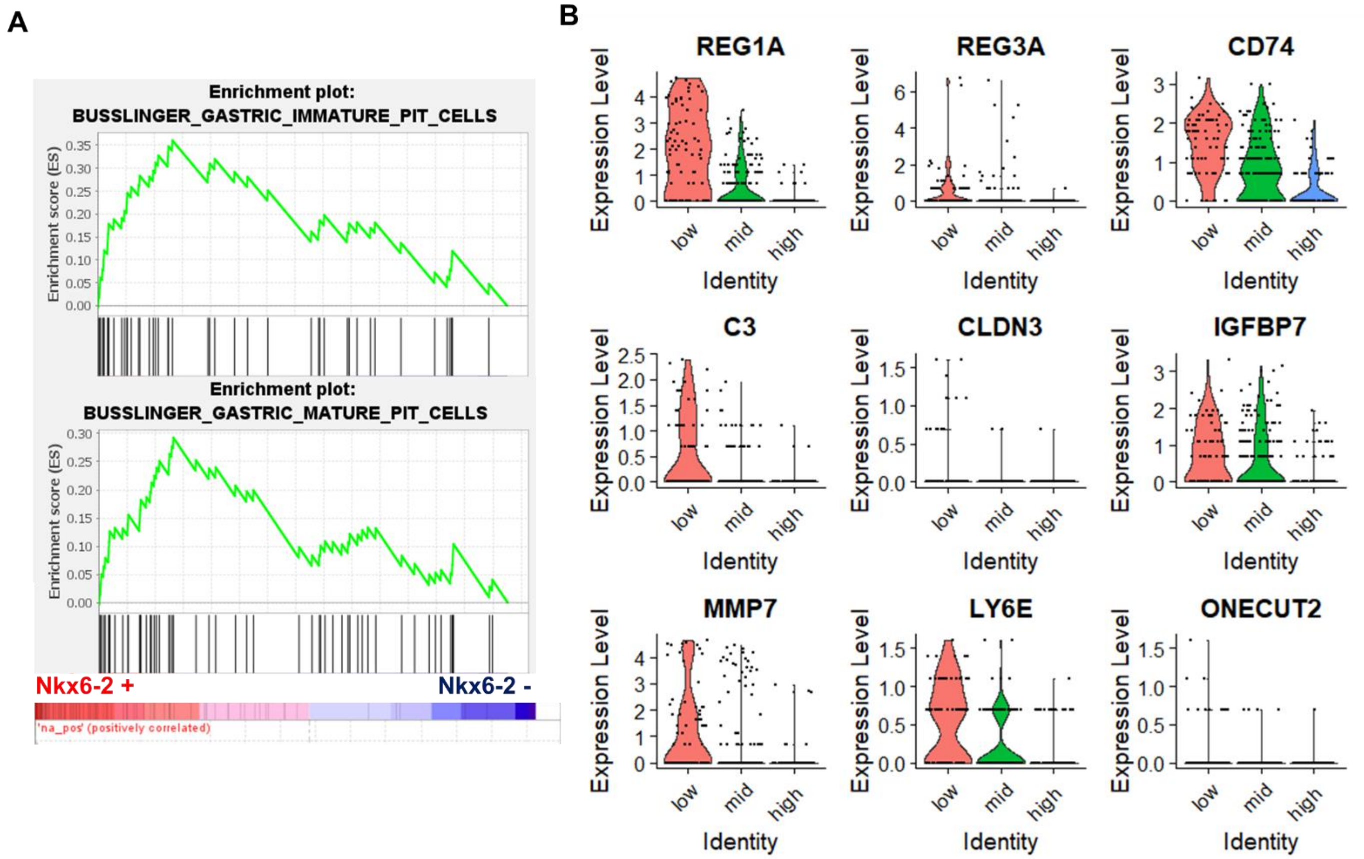


**Figure S10**. A) Gene set enrichment analyses (GSEA) of differentially expressed cell type signatures between Nkx6-2+ and Nkx6-2 – spots in the *Gnas;Kras* mice samples. Cell type signature enrichment was performed using the cell type human gene set. B) Expression of gene transcripts in the human LG IPMN epilesional areas found at increased expression in spots with low *Nkx6-2* expression in the Kras;Gnas murine ST dataset showing overlap in the trends observed in both datasets.

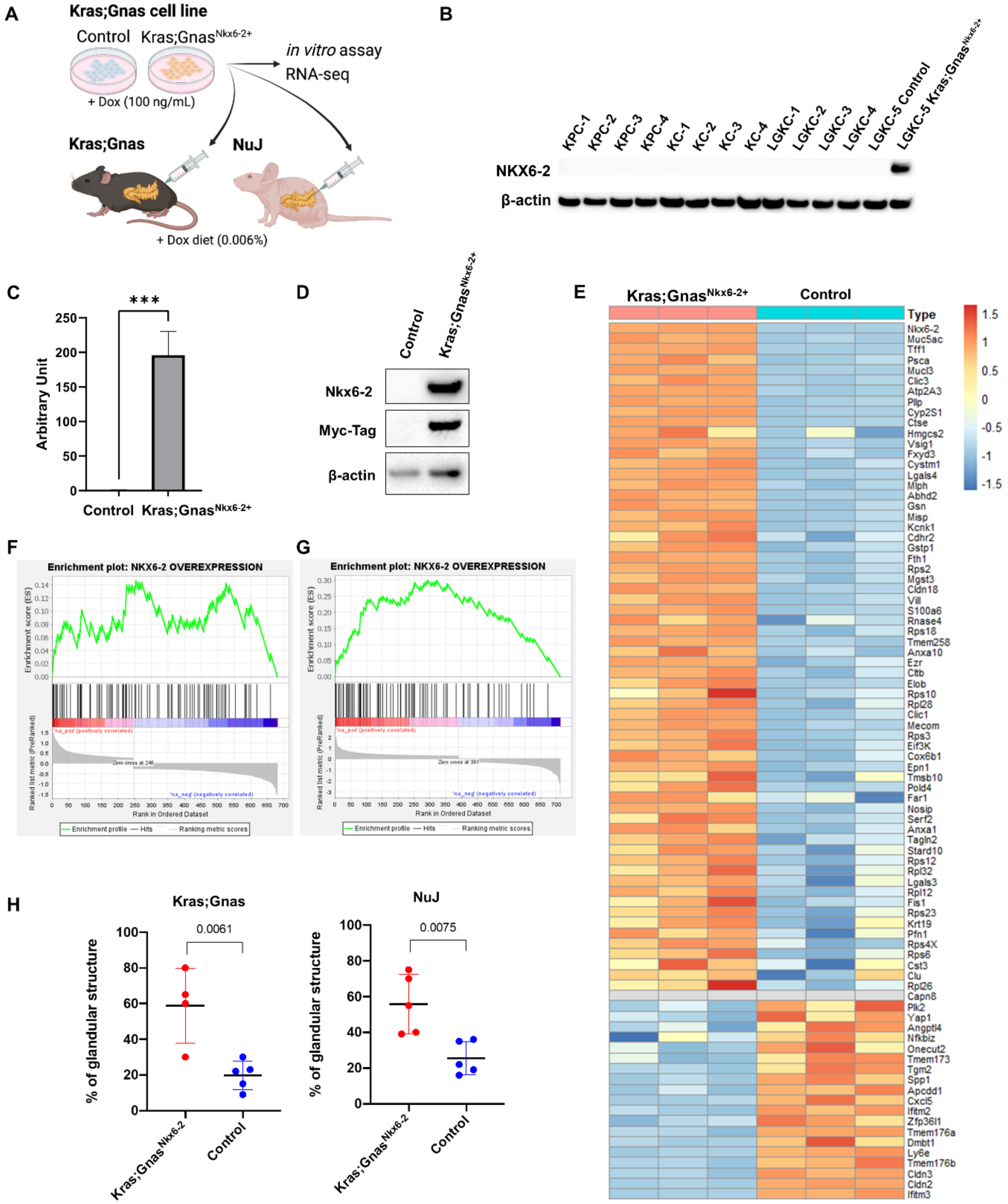


**Figure S11**. **Nkx6-2 overexpression in a Kras;Gnas cell line.** A) Experimental scheme of *in vitro* assay, RNA sequencing and orthotopic injection using the control and Kras;Gnas$^{Nkx6\text{-}2+}$ cells created with Biorender.com. B) Western blot of cell lysates obtained from four K-ras$^{LSL.G12D/+}$; and Trp53$^{R172H/+}$; Pdx-1-Cre (KPC), four K-ras$^{LSL.G12D/+}$; Pdx-1-Cre (KC) and four Kras;Gnas (LGKC) cell lines, stained with NKX6-2 and β-actin antibodies. The control and Kras;Gnas$^{Nkx6\text{-}2+}$ cell lines were used as a negative or positive control for NKX6-2 expression, respectively. C) Expression of Nkx6-2 in a cell line derived from Kras;Gnas mice transduced with either the control or Nkx6-2 vector (Kras;Gnas$^{Nkx6\text{-}2+}$) determined by qPCR. *** $P < 0.001$. D) Western blot of cell lysates obtained from the control and Kras;Gnas$^{Nkx6\text{-}2+}$cell line, stained with NKX6-2, Myc-Tag, and β-actin antibodies. E) Heatmap showing normalized expression levels of genes found to correlate with *Nkx6-2* expression in both the Kras;Gnas cell line and the LG IPMN ST dataset. F-G) Enrichment of the *Nkx6-2* overexpression gene set in the gene profiles obtained from differential expression of *Nkx6-2* + vs *Nkx6-2* – spots in the Kras;Gnas mice (NES = 1.48, FDR q-val = 0.09, D) and human LG IPMN tissues (NES = 2.93, FDR q-val = 0. H) Quantification of the percentage of glandular structures in the tumors derived from Kras;Gnas$^{Nkx6-2}$ or control cell lines in littermate (Kras;Gnas, left) mice or immunodeficient nude mice (NuJ, right).

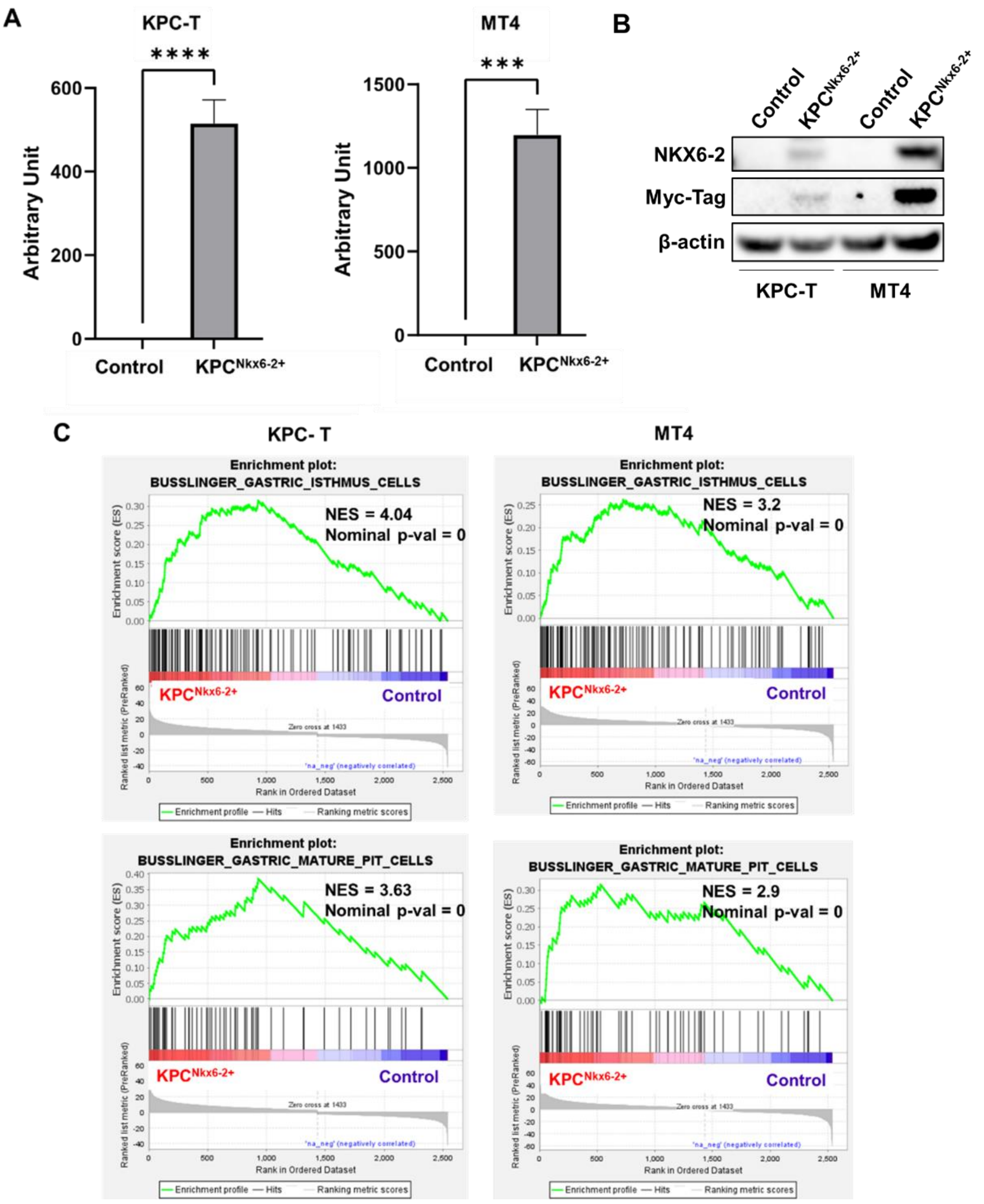


**Figure S12. Nkx6-2 overexpression in KPC cell lines.** A) Expression of Nkx6-2 in two KPC cell lines derived from K-ras$^{LSL.G12D/+}$; and Trp53$^{R172H/+}$; Pdx-1-Cre mice transduced with either the control or Nkx6-2 vector (KPC$^{Nkx6-2+}$) determined by qPCR. *** $P < 0.001$, **** $P < 0.0001$. B) Western blot of cell lysates obtained from the control and KPC$^{Nkx6-2+}$ cell line, stained with NKX6-2, My-Tag, and β-actin antibodies. C) Gene set enrichment analyses (GSEA) of differentially expressed genes between KPC$^{Nkx6-2+}$ and control cell lines showing enrichment of gastric isthmus and gastric pit gene sets.

### Supporting Methods

### <u>COMET Cyclic IF Staining</u>

#### Experimental workflow

The Lunaphore COMET (Switzerland) was employed to perform staining and imaging on the FFPE tissue samples. In brief, FFPE slides were placed in tanks containing BioGenex (CA, USA) EZ Elegans AR 2 buffer, which was in turn enclosed inside a BioGenex EZ Retriever microwave system. The samples were microwave heated to 107$^{o}$C for 15 min to dewax, rehydrate, and retrieve the antigens. Upon completion of heating the slides were cooled to room temperature and then loaded into the COMET and covered and sealed by a Fast-Fluidic Exchange microfluidics chip for staining and imaging in the instrument. The COMET performs a sequential immunofluorescence-based staining and imaging followed by an elution of the primary and secondary antibodies. Antibodies were titrated for optimal signal and verified to elute from the tissue using a "Characterization Type 2" instrument protocol where slides are stained, imaged, eluted, re-imaged for elution verification, and then subsequently re-stained etc. During the titration steps these elution images were collected to demonstrate the system performed as prescribed and titrations were empirically determined. During data collection staining and imaging elution images were not collected, thus the imager would stain, image, and elute; repeating these cycl es 10 times (2 primary Ab and 2 secondary Ab per cycle) to collect the information on the 20 Ab targets used in this study. The COMET collects images in OME-Tiff formats which were then visualized in COMET Viewer (Lunaphore, Switzerland).

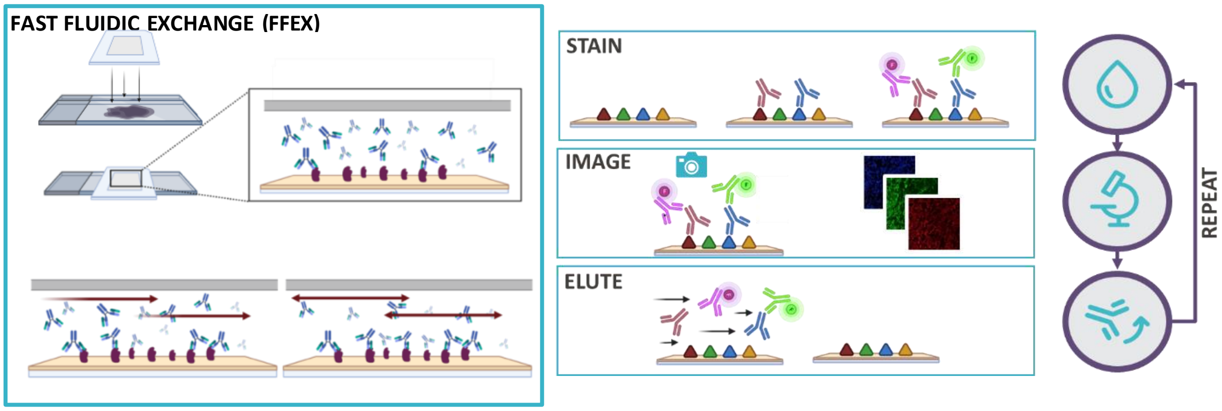


Staining was performed for 4-8 minutes per cycle for primary anitbodies which were diluted 1:50-1:250 in Multistaining Buffer (Lunaphore, Switzerland). Seconary antibodies, Alexafluor 555 and Alexafluor 647 (ThermoFisher) were diluted 1:200 and 1:400 respectively in Multistaining Buffer. Tissues were counterstained with DAPI solution (ThermoFisher) at every cycle. Imaging was performed on the COMET at 80 msec for DAPI, 400 msec for Alexafluor 555, and 200 msec for Alexafluor 647 at each cycle. Elution was peformed for 2 min using Elution Buffer (Lunaphore, Switzerland).

#### Analysis

Images were imported into the Visiopharm software. The epithelium of IPMN regions in the images was outlined and nuclei were segmented. The adjacent microenvironment areas were selected and cells were segmented. Expression of NKX6-2 and GATA-6 was measured per nuclei, and MUC5AC, TFF1, and VIM expression was quantified per sample. For phenotyping of the cells in the adjacent microenviroment, the order shown in Table S7 was followed. Intensities for all channels were background substracted using the individual signals from the secondary antibodies. Intensity cut-offs for determining positive and negative signals were done by evaluating several images collected in different runs and by comparing their values compared to the background.

### Immunofluorescence staining

Kras;Gnas mouse tissues containing LG IPMN, HG IPMN and PDAC were obtained at 18, 28 or later weeks on doxycycline diet (13). Five micrometer tissue sections were de-paraffinized in Xylene and rehydrated through 100%, 95% and 75% ethanol. Antigen retrieval was performed using a pH 6 citrate buffer (Dako) in a microwave for 15 min at 98°C. Each section was incubated in a protein block solution (Dako) for 1.5 hours at room temperature and incubated in NKX6-2 (Affinity DF9590, 1:100) and P120 (Proteintech 66208-1-IG, 1:500) primary antibodies overnight at 4°C. After three washes with TBS, the sections were incubated in donkey anti-rabbit IgG Alexa Fluor 488 (ThermoFisher A-21206, 1:500) and donkey anti-mouse IgG Alexa Fluor 647 (ThermoFisher A-31571, 1:500) for 1 hour at room temperature and DAPI diluted in PBS (1:5000) was applied to each section for 3 min for a nuclei counterstain. Slides were mounted with ProLongTM Diamond Antifade Mountant (ThermoFisher P36970) and cover slipped. Images were captured using an Olympus FV1000 laser scanning microscope.

### Western blot

To obtain protein extracts, cell pellets were lysed by RIPA Lysis Buffer (Rockland Immunochemicals, Philadelphia, PA, USA) containing protease/phosphatase inhibitor cocktail. Electrophoresis was performed to separate proteins in 4%–12% of Bis-Tris gels (Thermo Fisher Scientific, Waltham, MA, USA). The samples were subsequently transferred onto a nitrocellulose membrane. After blocking by 5% dry milk, membranes were incubated with anti-NKX6.2 (1:1000 dilution) (Millipore, Billerica, MA), anti-Myc-Tag (1:1000) (Cell Signaling Technology, Beverly, MA, USA), anti-E-cadherin (1:1000) (Cell Signaling Technology, Beverly, MA, USA), anti-Vimentin (1:1000) (Cell Signaling Technology, Beverly, MA, USA), anti-ZEB1 (1:2000) (Novus Biologicals, Littleton CO, USA), anti-ZEB2 (1:1000) (Invitrogen, Carlsbad, CA, USA) or HRP-conjugated anti-β-actin (1:10,000) (Santa Cruz Biotechnology, Dallas, TX, USA) primary antibodies at 4°C overnight. Blots for NKX6.2 and Myc-Tag were thereafter incubated with an HRP-conjugated anti-rabbit secondary antibody (Cell Signaling Technology). Clarity ECL Western Blotting Substrates (Bio-Rad, Hercules, CA) were used to visualize the blots. For RNA collection, GFP-positive cells were sorted from the cells transduced with a Nkx6-2 vector or a control vector using a CytoFLEX SRT (Beckman Coulter Life Sciences, CA, USA). 24 hours after plating, cells were treated with 100 ng/mL doxycycline for 24 hours before RNA collection.

### Quantitative real-time PCR and RNA sequencing

Total RNA was extracted using the RNeasy Mini Kit (QIAGEN, Venlo, Limburg, Netherlands), according to the manufacturer's protocol. cDNA was subsequently produced by reverse transcription. Gene expression was measured by real-time RT-PCR using the delta-delta Ct method. Following primers for PCR were purchased from Integrated DNA Technologies (IDT) (Coralville, IA, US); mouse NKX6-2 forward (5'-AAGTCTGCCCCGTCTCAAC-3'), mouse NKX6-2 reverse (5'-GGTCTGCTCGAAAGTCTTCTC-3'), mouse actin forward (5'-CACAGCTTCTTTGCAGCTCCTT-3'), and mouse actin reverse (5'-CGTCATCCATGGCGAACTG-3'). Bulk RNA sequencing was performed in triplicate on a NextSeq500 (Illumina, CA, USA). Sequencing reads were mapped using STAR (1) against the Mus musculus genome (UCSC mm10), and gene level quantification was performed with featureCounts (2) using the mm10 genome transcriptome annotation. Differential expression analysis was performed with DESeq2 (3).
